# Discovery of a novel merbecovirus DNA clone contaminating agricultural rice sequencing datasets from Wuhan, China

**DOI:** 10.1101/2023.02.12.528210

**Authors:** Adrian Jones, Daoyu Zhang, Steven E. Massey, Yuri Deigin, Louis R. Nemzer, Steven C. Quay

## Abstract

HKU4-related coronaviruses are a group of betacoronaviruses belonging to the same merbecovirus subgenus as Middle Eastern Respiratory Syndrome coronavirus (MERS-CoV), which causes severe respiratory illness in humans with a mortality rate of over 30%. The high genetic similarity between HKU4-related coronaviruses and MERS-CoV makes them an attractive subject of research for modeling potential zoonotic spillover scenarios. In this study, we identify a novel coronavirus contaminating agricultural rice RNA sequencing datasets from Wuhan, China. The datasets were generated by the Huazhong Agricultural University in early 2020. We were able to assemble the complete viral genome sequence, which revealed that it is a novel HKU4-related merbecovirus. The assembled genome is 98.38% identical to the closest known full genome sequence, *Tylonycteris pachypus* bat isolate BtTp-GX2012. Using *in silico* modeling, we identified that the novel HKU4-related coronavirus spike protein likely binds to human dipeptidyl peptidase 4 (DPP4), the receptor used by MERS-CoV. We further identified that the novel HKU4-related coronavirus genome has been inserted into a bacterial artificial chromosome in a format consistent with previously published coronavirus infectious clones. Additionally, we have found a near complete read coverage of the spike gene of the MERS-CoV reference strain HCoV-EMC/2012, and identify the likely presence of a HKU4-related-MERS chimera in the datasets. Our findings contribute to the knowledge of HKU4-related coronaviruses and document the use of a previously unpublished HKU4 reverse genetics system in apparent MERS-CoV related gain-of-function research. Our study also emphasizes the importance of improved biosafety protocols in sequencing centers and coronavirus research facilities.

## Introduction

Coronaviruses (CoVs) are a large group of RNA viruses infecting a range of animals, including humans. The Severe Acute Respiratory Syndrome Coronavirus (SARS-CoV) epidemic in 2002-2003, the Middle Eastern Respiratory Syndrome Coronavirus (MERS-CoV) epidemic in 2012-2015 with sporadic outbreaks through 2022 (World Health Organization, 2022) and the SARS-CoV-2 pandemic which began in late 2019 highlight the potential pathogenicity of coronaviruses to humans. MERS-CoV was first identified in 2012 (Bermingham et al., 2012), with 2,600 cases confirmed as of October 2022, and a case fatality rate of 36% (World Health Organization, 2022). The source of the initial MERS-CoV human infection was traced to dromedary camels on the Arabian Peninsula (Alraddadi et al., 2016; El-Kafrawy et al., 2019). Camels, however, are likely only intermediate hosts, with insectivorous bats the likely ancestral hosts of MERS-CoV (Lau et al. 2018).

MERS-CoV belongs to the merbecovirus (previously called lineage C) subgenus of the betacoronavirus genus of coronaviruses. Within merbecoviruses, in addition to the MERS-related CoV group, there are two other main phylogenetic groupings: HKU4-related CoVs and HKU5-related CoVs. HKU4-related CoVs were first identified in *Tylonycteris pachypus* bats in the Hong Kong Special Administrative Region (Woo et al., 2006), and since have been identified in *Tylonycteris* spp. across Southern China (Tang et al., 2006, Wu et al., 2016, Li et al., 2020). Notably, only a single HKU4-related CoV, Ty-BatCoV HKU4 SM3A, has been documented as having been isolated, with the virus replicating efficiently in human Caco-2 and Huh7 cells (Lau et al., 2021). HKU5-related CoVs are hosted in *Pipistrellus abramus* bats also found in Southern China (Woo et al., 2006).

The genomes of coronaviruses contain four genes coding for structural proteins: spike (S), envelope (E), membrane (M), and nucleocapsid (N). The coronavirus genome also contains a large replicase gene encoding (orf1ab), 5’ leader and untranslated region (UTR) and 3′ UTR and poly (A) tail (Fehr and Perlman, 2015). The S protein is responsible for binding to a host cell receptor on the cell surface and facilitating host cell entry.

In coronaviruses, the trimeric S protein consists of S1 and S2 subunits in each monomer. In MERS-CoV, SARS-CoV, and HKU4-related CoVs, the S1 subunit consists of an N-terminal domain and receptor-binding domain (RBD). The RBD of MERS-CoV binds to the human dipeptidyl peptidase 4 (hDPP4) receptor, which allows MERS-CoV to infect host cells (Raj et al., 2013). Although the HKU4-CoV S protein RBD also binds to hDPP4, its binding affinity is less than that of MERS-CoV (Yang et al., 2014). Notably however HKU4-CoV has been demonstrated to infect hDPP4 transgenic mice (Lau et al., 2021). HKU5r-CoVs however, lack the ability to bind to hDPP4 (Yang et al., 2014).

After the binding of the RBD with hDPP4, MERS-CoV relies on host cell protease cleavage to activate membrane fusion and gain cell entry (Millet and Whittaker 2014; Kleine-Weber et al., 2018). Proteolytic cleavage occurs at two positions within the S protein, at both the boundary of the S1 and S2 subunits (S1/S2), and adjacent to the fusion peptide within the S2 subunit (S2′) (Yamada and Liu 2009; Belouzard et al., 2009; Fehr and Perlman 2015). Unlike MERS-CoV, HKU4-related CoVs do not possess furin cleavage motifs at either the S1/S2 boundary or the S2’ location, and cannot efficiently utilize endogenous human proteases for cell entry (Yang et al., 2014; Wang et al., 2014; Yang et al., 2015). However Yang et al. (2015) demonstrated that by introducing two mutations at the S1/S2 boundary of HKU4-CoV to produce a furin cleavage site, the HKU4-CoV S protein was able to mediate entry into human cells.

Sequence Read Archive (SRA) data submitted to The National Center for Biotechnology Information (NCBI) may contain contamination, either through library or sample contamination prior to sequencing, cross-contamination between samples, or index-hopping within multiplexed runs (Ballenghien et al., 2017). Indeed, Katz et al. (2021) identified that in early 2020, over 2,000 Public Health England surveillance bacterial next-generation sequencing SRA datasets were likely contaminated with SARS-CoV-2 sequences. In some cases, the identification and characterization of cross-contaminating reads may provide useful biosecurity and or research insights (Lewis et al., 2020; Csabai and Stéger 2021; Quay et al., 2021; Jones et al., 2022).

BioProject PRJNA602160 on NCBI contains 26 SRA datasets containing *Oryza sativa* subsp. *japonica* (Japonica rice) DNA and RNA sequencing data (Supp. Info. 1.1). We identified several SRA datasets in this BioProject that contain HKU4-related CoV and MERS-related CoV sequences during a routine search of agricultural sequencing BioProjects from China. The datasets were generated by the Huazhong Agricultural University (HZAU), registered on 2020-01-19 and published on NCBI on 2020-02-09. BioProject PRJNA602160 utilized Bisulphite-sequencing and RNA-sequencing to characterize DNA methylation patterns and analysis of DNA glycosylases functionality in rice eggs, sperm cells, and zygotes. As a rice sequencing study, only rice genomic sequences, rice crop hosted viruses, and bacterial sequences are expected to be present in the SRA datasets.

Using a bioinformatics workflow that included *de novo* assembly, as well as read and contig alignments, we identified a novel HKU4-related CoV genome sequence in four SRA datasets in rice sequencing BioProject PRJNA602160. We show that the novel HKU4-related CoV likely binds to hDPP4, potentially representing a human spillover risk. Additionally, we unexpectedly identified that the coronavirus genome was contained in a bacterial artificial chromosome (BAC) plasmid. This represents the first reverse genetics system documented for HKU4-related CoVs. In addition to the identification of a novel HKU4-related CoV in a BAC, we identified a near complete MERS-CoV spike sequence. We show the MERS-CoV spike was very likely substituted into the novel HKU4-related CoV backbone, representing a second clone in the datasets. Such research is indicative of enhanced potential pandemic pathogen (gain-of-function) research and we assess how this novel HKU4r-related clone may have contaminated agricultural rice sequencing datasets.

## Results

### Identification of a novel merbecovirus

Of the 26 SRA datasets in BioProject PRJNA602160, *Tylonycteris* sp. bat CoV HKU4 sequence matches were identified in four datasets using the NCBI SRA Taxonomy Analysis Tool (STAT) (Katz et al., 2021), as tabulated in Supp. Info. 1.2. *Tylonycteris* bat CoV isolate BtTp-GX2012 was found to be the HKU4 strain with the highest percentage of matching reads in SRA datasets SRR10915168 and SRR10915173-4. To confirm the presence of HKU4-related CoV sequences in the four SRA datasets we undertook genome alignment analysis as described in the Methods section.

To extract a complete genome sequence, *de novo* assembly of each of the four SRA datasets in BioProject PRJNA602160 containing HKU4-related CoV sequences was conducted using MEGAHIT (Li et al., 2015). The resulting contigs were aligned to a combined set of the complete NCBI viral database and all coronaviruses on NCBI using minimap2 (Li, 2018). Of the 40 contigs matching HKU4-related CoVs or MERS-CoV, contig k141_13282 from assembly of SRA dataset SRR10915173 was found to have 100% coverage of the BtTp-BetaCoV/GX2012 genome. This contig was queried against the NCBI nucleotide (nt) database (Johnson et al., 2008) using NCBI BLAST (blastn), and the highest identity match was found to be *Tylonycteris* bat coronavirus isolate BtTp-GX2012 (Supp. Info. 1.3). The four bat coronavirus genomes with highest identity to contig k141_13282 were then input as queries against contig k141_13282 as a subject using blastn. The overall identity of the closest known genomes to the novel HKU4-related CoV was between 97.97% and 98.38% (Table 1). We named the novel HKU4-titlerelated CoV ‘HKU4r-HZAU-2020’ to reflect its phylogenetic affiliation, institutional source, and date of sequencing.

**Table 1.** Blastn nucleotide sequence identity of the four most closely related sequences on GenBank to HKU4r-HZAU-2020.

| Description | Accession | Max Score | Coverage | % identity |
| --- | --- | --- | --- | --- |
| BtTp-BetaCoV/GX2012 | KJ473822.1 | 53149 | 100% | 98.38% |
| T. BtCov HKU4 isolate CZ07 | MH002338.1 | 52711 | 99% | 98.14% |
| T. BtCov HKU4 isolate CZ01 | MH002337.1 | 52436 | 99% | 97.97% |
| BtCoV/133/2005 | DQ648794.1 | 48710 | 99% | 95.67% |

A BLAST search of HKU4r-HZAU-2020 against all HKU4-related sequences was conducted against the nt database on NCBI. Critical components of HKU4r-HZAU-2020, including the spike glycoprotein and the membrane glycoprotein, bear substantial differences when compared with other known sequences of HKU4 (Table 2).

**Table 2:** Nucleotide percentage identity of HKU4r-HZAU-2020 to the four most closely related sequences on GenBank for each major gene sequence. See Supp. Info. 1.4 for Genbank accession numbers.

| ORF1ab | % id | S | % id | E | % id | M | % id | N | % id |
| --- | --- | --- | --- | --- | --- | --- | --- | --- | --- |
| GX2012 | 98.86 | HKU4 CZ07 | 97.59 | HKU4 CZ07 | 99.6 | GX2012 | 98.33 | HKU4 CZ07 | 99.06 |
| HKU4 CZ07 | 98.61 | GX2012 | 96.62 | GX2012 | 99.6 | GZ131656 | 97.88 | HKU4 CZ01 | 99.06 |
| HKU4 CZ01 | 98.53 | HKU4 CZ01 | 96.53 | HKU4 CZ01 | 99.2 | 133/2005 | 97.73 | GX2012 | 99.06 |
| BtCoV/133/2005 | 95.99 | GZ160421 | 96.23 | GZ131656 | 98.8 | HKU4-4 | 96.36 | 133/2005 | 97.18 |

### Characterization of HKU4r-HZAU-2020 as a cDNA clone

The length of contig k141_13282 at 38,583 nt was significantly longer than HKU4-related or MERS-related CoV genomes, which are 30 kb to 33 kb in length. Reads in each each of the four SRAs containing HKU4r-HZAU-2020 sequences were aligned to contig k141_13282, and the regions covering the 5’ and 3’ ends of the HKU4r-HZAU-2020 genome were analyzed using Addgene sequence analyser (Addgene, 2022), Integrative Genomics Viewer (IGV) (Thorvaldsdóttir et al., 2013), and blastn. A human cytomegalovirus (CMV) immediate-early promoter sequence was identified immediately upstream of the 5’ end of the HKU4r-HZAU-2020 genome (Supp. Fig. 1). Trailing the poly(A) tail at the 3’ end of the genome, a hepatitis delta virus (HDV) ribozyme and bovine growth hormone (bGH) polyadenylation signal was found (Supp. Fig. 2). With a read depth of 40-60 over 200 nt regions centered on both the 3’ and 5’ junction locations, we infer that synthetic sequences were attached to the genome, with good confidence.

The first 316 nt of the 6343 nt sequence attached to the poly(A) tail at the 3’ end of the HKU4r-HZAU-2020 genome exhibited the highest blastn maximum score match to vector pBAC-Beaudette-FU (MW847254.1) (92.11% identity, 97% coverage), which is a BAC plasmid containing infectious bronchitis virus (Inayoshi et al., 2022). The trailing 6027 nt was identified using blastn as having a 100% identity match to the BAC cloning vector pDEV-CHa (MT702985.1) (Chen et al., 2013). Overall however, the highest blast maximum score match we identified to sequences on the NCBI nt database was found using the *align two or more sequences* option to Sequence 11 from Patent WO2006236448 “Attenuated SARS and use as a vaccine” (CS480537.1) (Enjuanes et al. 2006) with a 6329/6368 nt (99.39%) match (Supp. Info. 1.5). The matching section of Sequence 11 from Patent WO2006236448 forms part of a pBAC-SARS-CoV vector backbone (Enjuanes et al., 2006; Almazán et al., 2006; Almazán et al., 2008). The first 627 nt of the HKU4r-HZAU-2020 sequence trailing the poly(A) tail of the genome has a similar HDV ribozyme and bGH polyadenylation signal layout to the pBAC-SARS-CoV Sequence 11 trailing a SARS poly(A) tail (Supp. Fig. 3).

A 1912 nt section of contig k141_13282 was found upstream of the 5’ end of the HKU4r-HZAU-2020 genome. Anomalous read depth and coverage was found over the first 149 nt of the sequence, and this region was trimmed from the contig. The resultant 1763 nt sequence was analyzed using blastn, with highest max score match to BAC constructs Gallid alphaherpesvirus 1 strain A489 (KY423284.1) and BAC cloning vector pDEV-CHa (MT702985.1). The sequence was compared with pBAC-SARS-CoV Sequence 11 using blastn and Addgene sequence analyser (Supp. Fig. 4), with two regions exhibiting a 100% identity, a 956 nt sequence includes a lambda cos and a loxP site, which are commonly used for phage package and cre cleavage, respectively (Wang et al., 2022). A 587 nt sequence includes a M13 forward primer and CMV promoter. The complete contig k141_13282 with 149 nt at the 5’ trimmed (‘k141_13282_del_149’) was annotated using Snapgene (Fig. 1). We refer to the recovered full length pBAC and genome sequence in contig k141_13282_del_149 as HKU4r-HZAU-2020, and refer to the viral genome sequence as the HKU4r-HZAU-2020 genome.

**Fig. 1.**
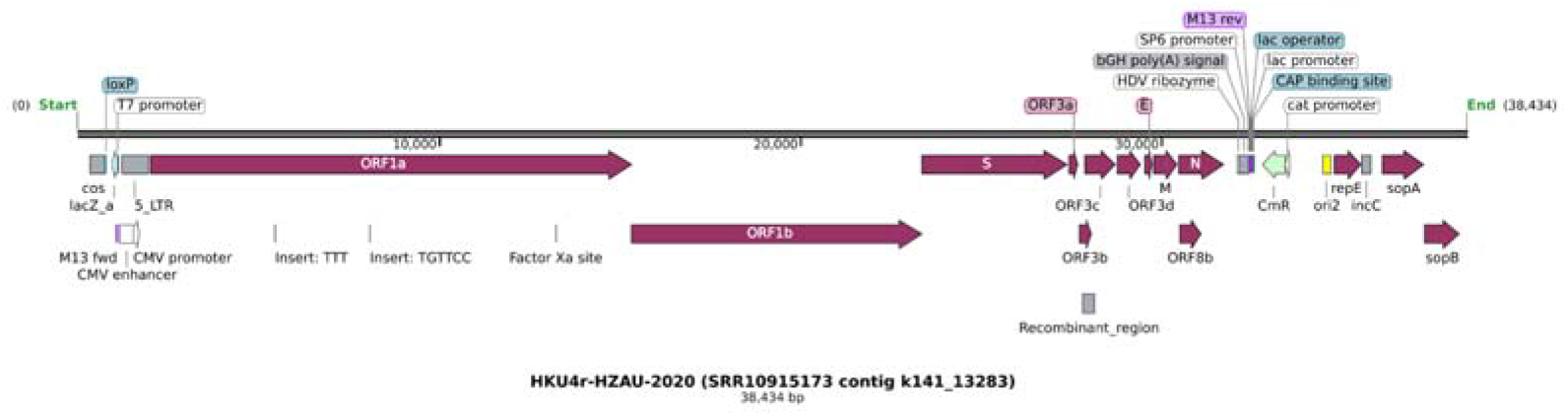
Annotation of the largest assembled HKU4-related contig sequence obtained from rice dataset SRR10915173. A 149 nt region at the 5’ end of the sequence with anomalously high coverage was removed prior to annotation.

The presence of T7 and CMV promoters before the 5’-end of the HKU4r-HZAU-2020 genome and a Hepatitis D virus ribozyme, followed by a bGH polyA signal after the 3’-end of the genome, indicates that the novel HKU4-related CoV obtained from SRR10915173 is probably infectious or intended to be infectious. This is evidenced by its format, which could generate full-length infectious RNA when expressed in mammalian cells.

### Quality control

Reads in each of the four SRAs in BioProject PRJNA602160 containing merbecovirus sequences were aligned using minimap2 to the full HKU4r-HZAU-2020 clone (38,434 nt trimmed contig ‘k141_13282_del_149’), as shown in Table 3. The four datasets were pooled and the combined reads again mapped to HKU4r-HZAU-2020 (Table 3).

**Table 3.** Number of reads, coverage and average read depth for each of the four SRA datasets (*and pooled datasets) containing merbecovirus sequences in BioProject PRJNA602160 mapping to the full HKU4r-HZAU-2020 sequence (trimmed contig ‘k141_13282_del_149’). Detailed statistics can be found in Supp. Info. 1.5.

| SRA | Reference | Average depth | Coverage% | Reads mapped |
| --- | --- | --- | --- | --- |
| 4 pooled SRAs* | HKU4r-HZAU-2020 | 65.6 | 100 | 19952 |
| SRR10915173 | HKU4r-HZAU-2020 | 59.68 | 100 | 18309 |
| SRR10915174 | HKU4r-HZAU-2020 | 3.6 | 93.68 | 1004 |
| SRR10915168 | HKU4r-HZAU-2020 | 1.91 | 77.41 | 518 |
| SRR10915167 | HKU4r-HZAU-2020 | 0.4 | 32.97 | 121 |

The process was repeated using two additional short read aligners, bowtie2 and bwa-mem2, with similar results. Minimap2 aligned read depth distribution was fairly consistent across the full contig sequence for the SRA dataset with highest HKU4-related CoV read count, SRR10915173, except for three local regions: positions 1-71 with read depth of 29-48; positions 1919-1973 with a read depth of 22-28 and located within the 5’ UTR sequence 155 nt downstream from the 5’ end of the HKU4r-HZAU-2020 viral genome; and between positions 32412 and 32521 immediately downstream of a bGH poly(A) signal and upstream of a CAP binding sequence, with a read depth between eight and 20 reads (Supp. Figs. 5-7). For the four pooled SRA datasets, slight differences in read depth were found between aligners betweens positions 32412 and 32521 of the HKU4r-HZAU-2020 clone with slightly higher read coverage over this section of the HKU4r-HZAU-2020 clone with minimap, bowtie2 and bwa-mem2 having minimum read depths of 9, 10 and 14 reads respectively (Supp. Figs. 8-10). While we infer that this read depth is sufficient to indicate the likely presence of a single contiguous sequence, we cannot conclusively rule out misalignment of a second vector sequence trailing the bGH poly(A) signal.

Since the choice of assembler can affect the quality of betacoronavirus *de novo* assembly (Islam et al., 2020), two other *de novo* assemblers were tested and the results compared with the MEGAHIT assembly. CoronaSPAdes (Meleshko et al., 2021) using default parameters and SPAdes (Prjibelski et al., 2020) with the ‘careful’ setting, were used to assemble each of the four SRA datasets containing merbecovirus sequences. Using SPAdes careful assembly of SRR10915173, a 38,592 nt contig with 100% coverage and identity to the HKU4r-HZAU-2020 complete sequence was recovered. At the same time, *de novo* assembly of SRR10915173 using coronaSPAdes generated two contigs of lengths 32,683 nt and 5879 nt, which had a combined 100% coverage and 100% identity to the HKU4r-HZAU-2020 complete sequence (Supp. Data).

To test the alignment to the HKU4r-HZAU-2020 viral genome sequence alone, each of the four SRAs with merbecovirus sequences, as well as a pooled set of these four datasets, were aligned to the HKU4r-HZAU-2020 viral genome sequence using minimap2, bowtie2 and bwa-mem2. In each of the four SRA datasets, at least one read crosses the 5’ end of the genome, with soft-clipped overhangs upstream of the 5’ end of the genome, all with a nearly identical sequence (Supp. Fig. 11). SRR10915173 has the most number of reads crossing the 5’ end of the genome with 38x 150 nt reads, while SRR10915174 has four 150 nt reads covering this region. As discussed above, a CMV forward primer sequence was found in the 5’ end of reads covering the region immediately upstream of the 5’ end of the HKU4r-HZAU-2020 genome. Similarly, a HDV ribozyme sequence was found in the 3’ end of reads covering the region immediately downstream of the poly(A) tail of the HKU4r-HZAU-2020 genome. This read configuration clearly demonstrates that the HKU4r-HZAU-2020 full genome had been assembled as part of a synthetic construct.

### Recombination and Simplot analysis

A recombination analysis using RDP4 (Martin et al., 2015), utilizing eight different methods, indicated only a single small 316 nt potential recombination fragment located between positions 26027 and 26343 in the HKU4r-HZAU-2020 genome (Fig. 1), which was detected by four methods (Supp. Info. 1.6). Thus, there is no clear evidence that the HKU4r-HZAU-2020 CoV identified here could have been the result of simple recombination from known HKU4 genomes.

Simplot analysis shows that three sequences: BtTp-BetaCoV/GX2012, HKU4 isolate CZ01 and HKU4 isolate CZ07 have high identity across the ORF1ab coding region, with CZ07 having the highest identity match in the S1 region of the spike gene (Fig. 2). The ORF3a/3b gene sections of the genome exhibit low identity to the HKU4-related CoV genomes analyzed and potentially represents a section of either natural or artificial recombination.

**Fig. 2.**
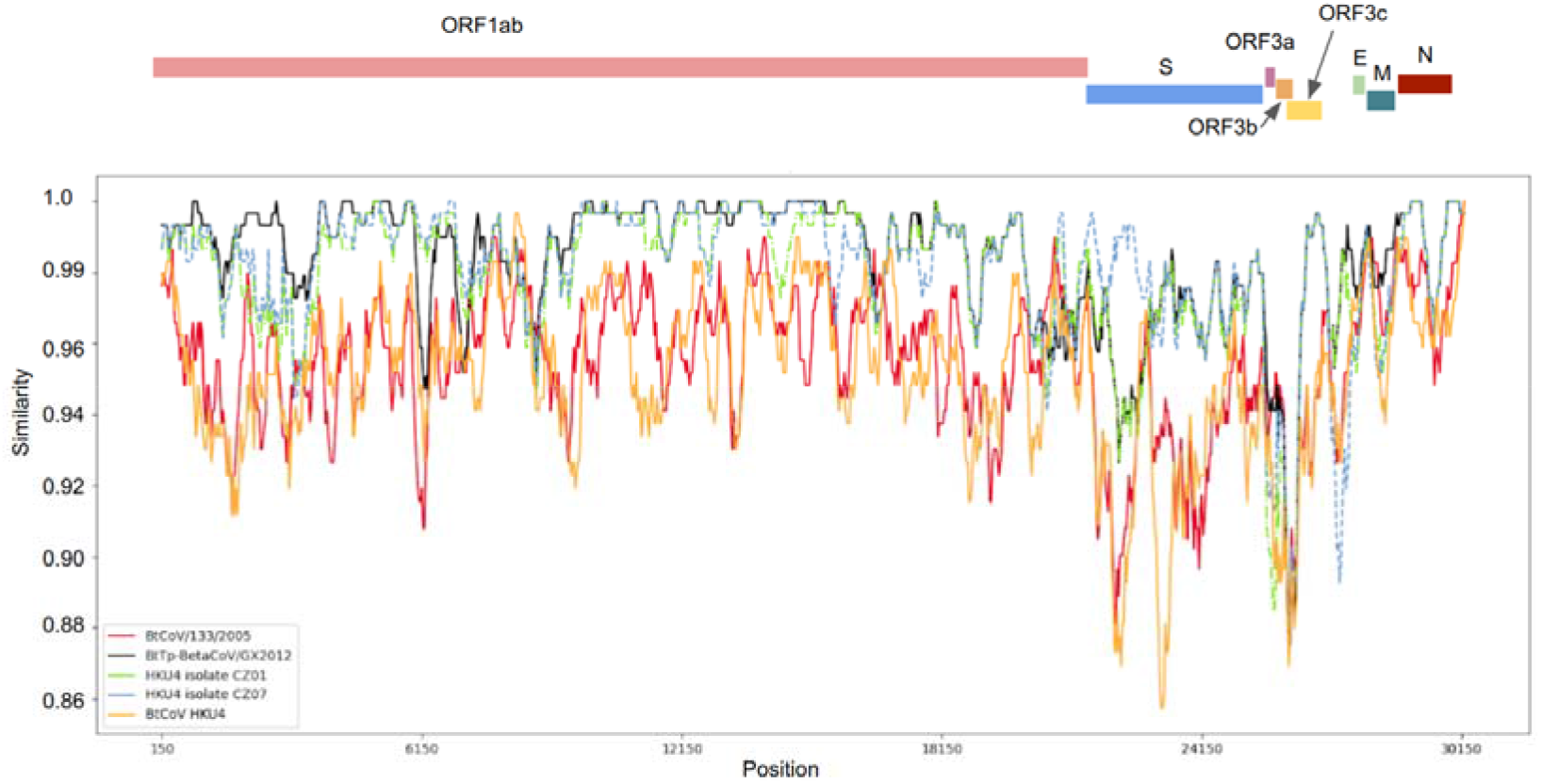
Similarity plot of the HKU4r-HZAU-2020 genome plotted against selected HKU4-related CoV genomes. HKU4 isolate CZ07 in light blue dashed pattern, HKU4 isolate CZ01 in green dash-dot pattern. Parameters: Window: 300 nt, step: 30 nt, model: Jukes-Cantor. Generated using Simplot++.

### Phylogenetic analysis

Maximum likelihood phylogenetic trees were generated for the full-length genome, spike protein coding sequence, RdRp gene, and partial RdRp gene. The HKU4r-HZAU-2020 genome forms a basal sister relationship to BtTp Beta-CoV/GX2012, HKU4 isolate CZ07 and HKU4 isolate CZ01 for the full genome (Fig. 3).

**Fig. 3.**
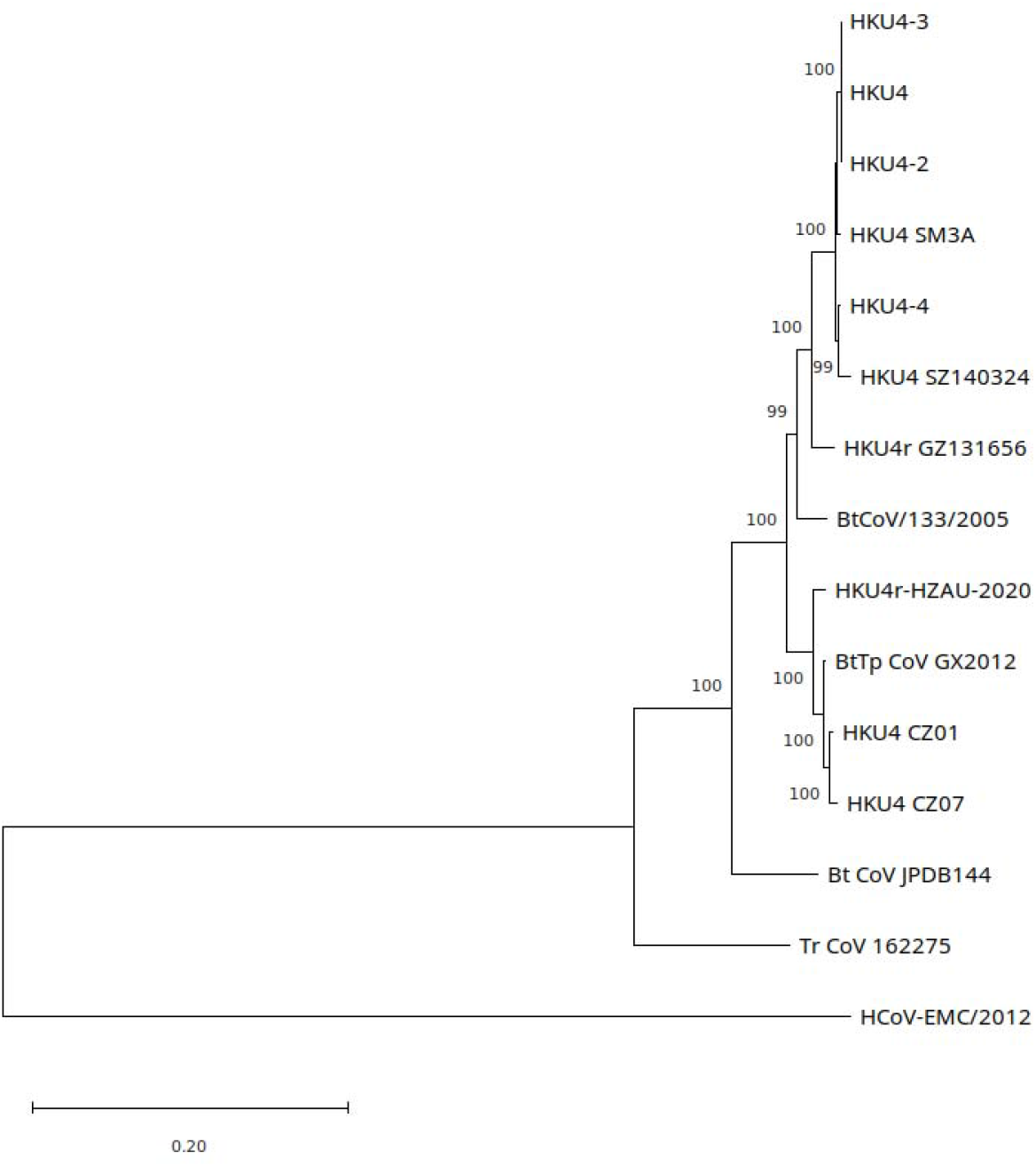
Maximum likelihood tree generated for selected HKU4-related CoV and MERS-CoV full genomes using a GTR +R model in PhyML using smart model selection (Lefort et al., 2017). Tree rooted on midpoint. See Supp. Info. 1.4 for Genbank accession numbers.

For the spike gene the HKU4r-HZAU-2020 genome forms a sister group to the HKU4 isolates CZ07/CZ01 and BtTp Beta-CoV/GX2012 clade. (Fig. 4). The sister clade to the HKU4r-HZAU-2020, HKU4 isolates CZ07/CZ01 and BtTp Beta-CoV/GX2012 clade consists of three *Tylonycteris pachypus* HKU4 isolates, GZ1912, GZ1862 and GZ1832. These were collected in Guizhou province, China on 2015-09-09 and sequences submitted to Genbank on 2020-11-04 by Lau et al. (2021).

Phylogenetic analysis of the S protein shows similar grouping of HKU4r-HZAU-2020 with the S proteins of HKU4 isolates CZ07/CZ01 and BtTp Beta-CoV/GX2012. The only documented HKU4 cell culture isolate, HKU4 SM3A, groups in a separate clade, with BtCoV/133/2005 and HKU4 in both S gene and S protein phylogenetic trees (Fig. 3; Supp. Fig. 12).

**Fig. 4.**
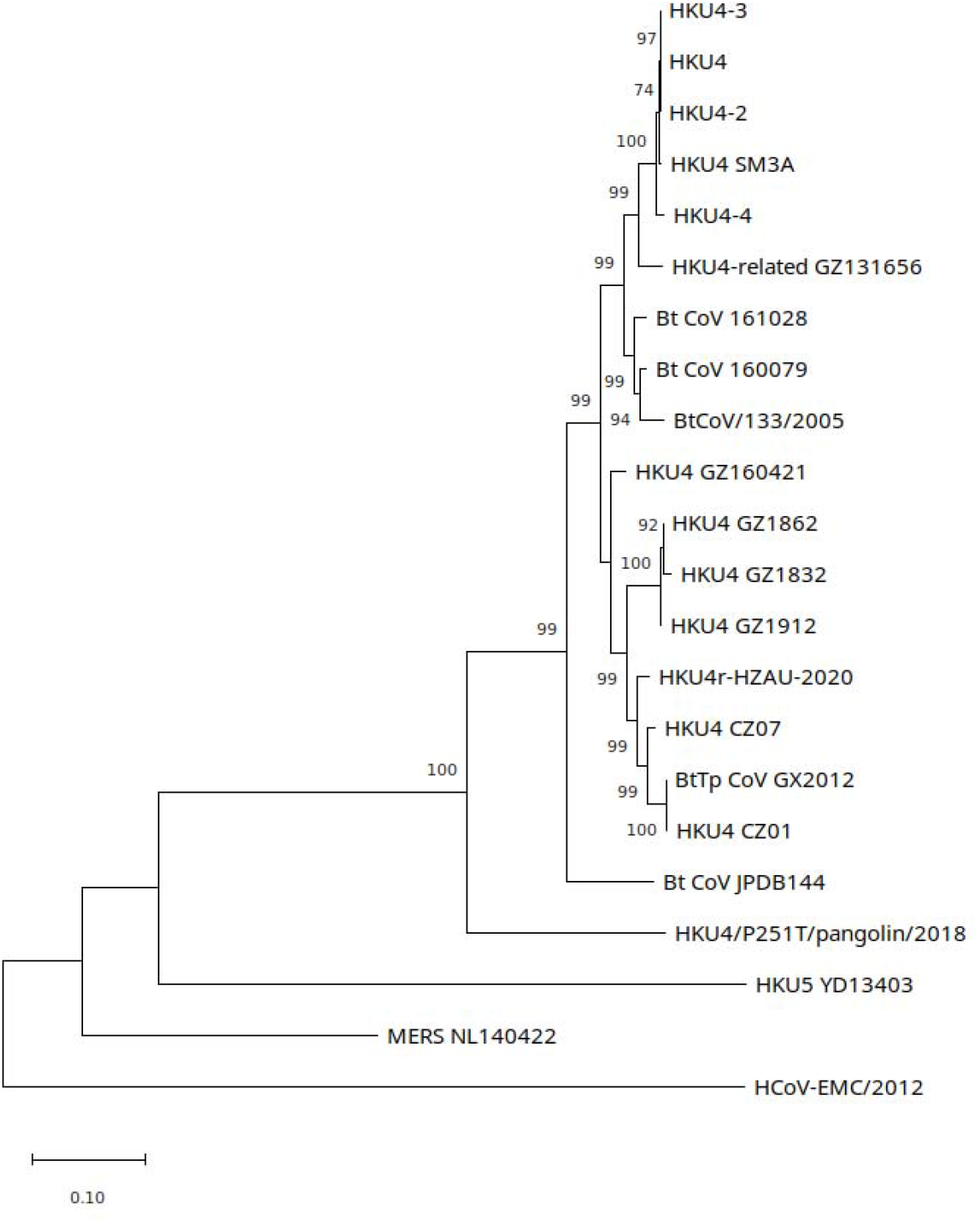
Maximum likelihood tree generated for HKU4-related CoV, MERS CoV and HKU5 CoV spike genes using a GTR +G model in PhyML using smart model selection (Lefort et al.). Tree rooted on midpoint. See Supp. Info. 1.4 for Genbank accession numbers.

For the RdRp gene, HKU4r-HZAU-2020, BtTp Beta-CoV/GX2012, HKU4 isolate CZ07 and HKU4 isolate CZ01 form an undifferentiated clade with good support (Fig. 5). Using a 399 nt section of the RdRp that was more widely sampled that the full RdRp, we identified the closest phylogenetic grouping to Tylonycteris bat coronavirus HKU4 isolate 152762 (MN312732.1), a *Tylonycteris pachypus* sequence collected from an undocumented location in China by Latinne et al. (2020) and submitted to Genbank on 2020-06-01 (Supp. Fig. 13).

**Fig. 5.**
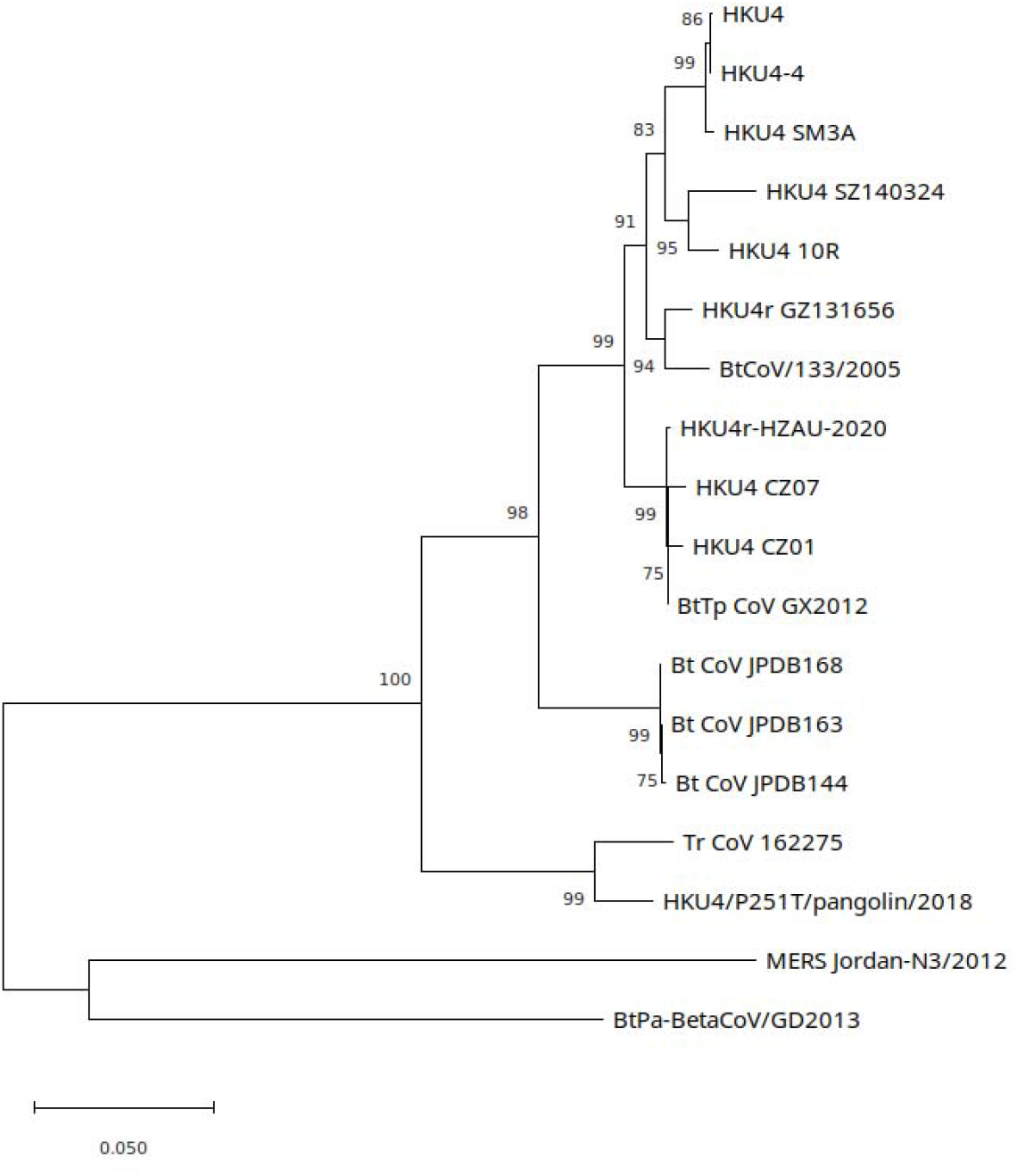
Maximum likelihood tree generated for selected HKU4-related CoVs, MERS-CoV and HKU5-related CoV RdRp genes using a GTR + G model in PhyML using smart model selection (Lefort et al., 2017). Tree rooted on midpoint. See Supp. Info. 1.4 for Genbank accession numbers.

We repeated phylogenetic analysis using the maximum likelihood method in MEGA11 with best fit model selection based on lowest Bayesian information criterion. 1,000 bootstrap replicates were calculated for full genome, S, and RdRp gene analysis. The respective topologies were consistent with those generated using PhyML (Supp. Figs. 14-16).

### HKU4r-HZAU-2020 host

In order to exclude the possibility that some of the sequences may have originated from a bat sample, we performed a blastn search using the available nucleotide sequences of the bat *Tylonycteris pachypus*, the reservoir host of HKU4, against SRR10915173. We did not obtain any sequence that matched any nucleotide sequence from this species.

In addition, we aligned each SRA dataset in BioProject PRJNA602160 against all mitochondrial genomes present on Genbank. We used a minimum genome coverage of 10% to infer presence (Supp. Fig. 17; Supp. Info. 2.1). Only *Danio rerio* (zebrafish), *Homo sapiens*, and three rice species *Oryza minuta*, *Oryza rufipogon*, and *Oryza sativa* had greater than 20% coverage, with several other plant, yeast, and parasites having between 10 and 20% mitochondrial genome coverage. For the four SRA datasets containing HKU4r-HZAU-2020 sequences, *H.sapiens* mitochondrial genome coverage varied between 37.1% and 38.4%. However, no bat mitochondrial genome coverage of greater than 0.8% was detected in any of the 26 SRA datasets in BioProject PRJNA602160.

We further identified rRNA matching reads in the four SRAs containing HKU4r-HZAU-2020 sequences SRR10915167-8 and SRR10915173-4 using Metaxa2. The reads were *de novo* assembled, and blastn was then used on the assembled contigs to identify closest identity in the NCBI nt database. *H.sapiens* mitochondrion rRNA matching contigs were found in all four datasets, comprising between 12.5% and 25% of the contigs (Supp. Info. 2.2; Supp. Data).

These results indicate that the HKU4r-HZAU-2020 clone was not associated with a bat host. Furthermore, the presence of vector sequences at the 5’ and 3’ ends of the genome indicate that it was not expressed as RNA, but rather it was present as DNA. This is also consistent with the constant read depth across the 5’ and 3’ vector/virus boundaries (Supp. Figs. 1-2, 7), and lack of marked variability in read depth across the genome sequence itself.

### Receptor binding domain in silico analysis

The RBD of HKU4-related CoVs are known to bind hDPP4 (Wang et al., 2014). We identified the closest match to the HKU4r-HZAU-2020 RBD protein sequence on the NCBI nr database as PDB structure 4QZV:B (Wang et al., 2014). To ascertain if the HKU4r-HZAU-2020 RBD was likely to also bind to hDPP4, we generated a structural model of the HKU4r-HZAU-2020 RBD using SWISS-MODEL and visualized its docking to human DPP4 using PyMol (Fig. 6). High structural homology between the RBD from the HKU4r-HZAU-2020 clone and the RBD of PDB structure 4QZV was observed.

**Fig. 6.**
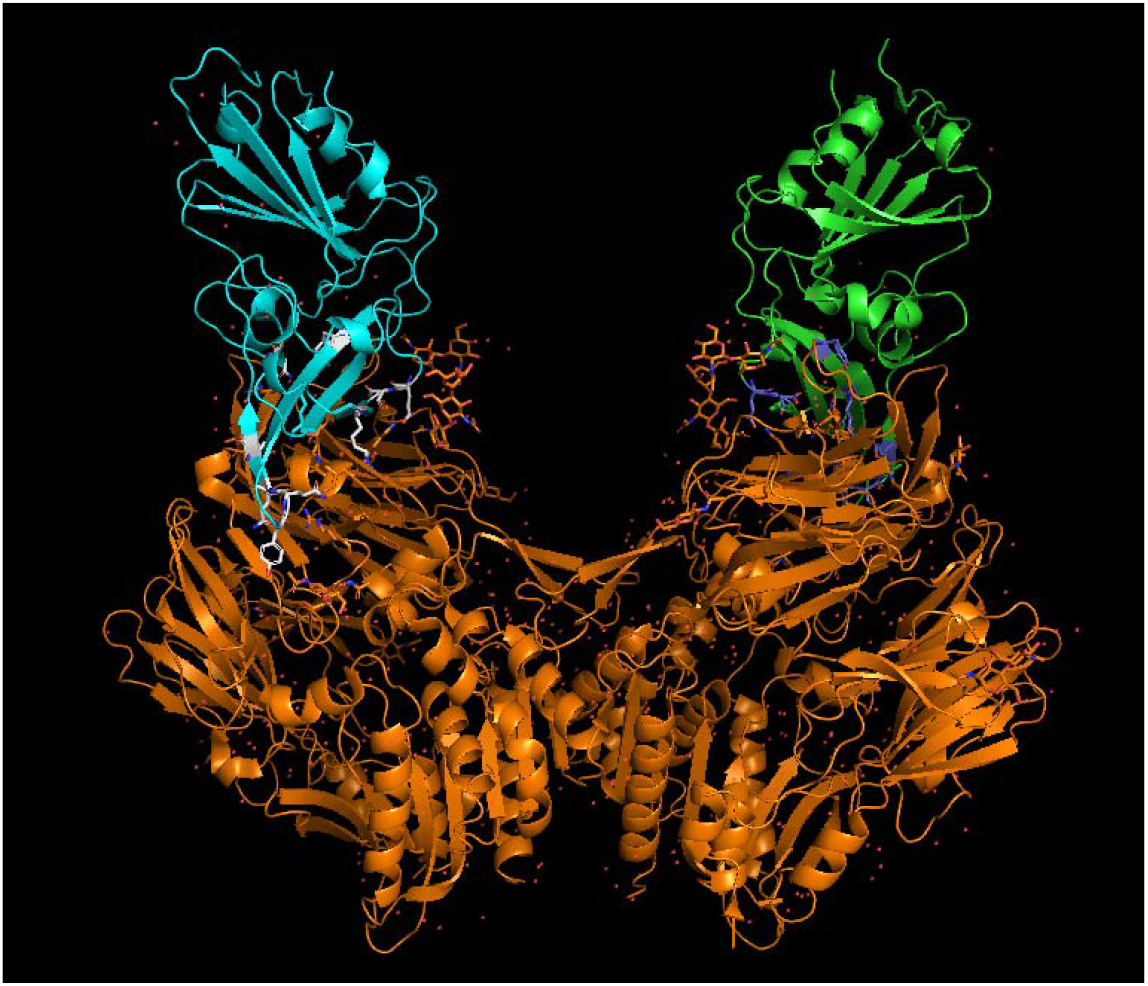
Alignment of the modeled RBD of the HKU4r-HZAU-2020 clone (light blue) to the HKU4 RBD in PDB (PDB ID 4QZV) (green) is consistent with HKU4r-HZAU-2020 RBD binding to hDPP4 (orange). Contacts between the RBD structures and hDPP4 were identified using PyMol (which were found to be 2.6 to 3.5 Angstroms in distance) and are indicated by blue sticks between the molecules.

We then undertook molecular docking modeling in PRODIGY, which showed comparable binding energy of the two RBD molecules to hDPP4 (Table 3), indicating that the HKU4r-HZAU-2020 CoV obtained from SRR10915173 may be capable of causing infection in humans.

**Table 3a.**
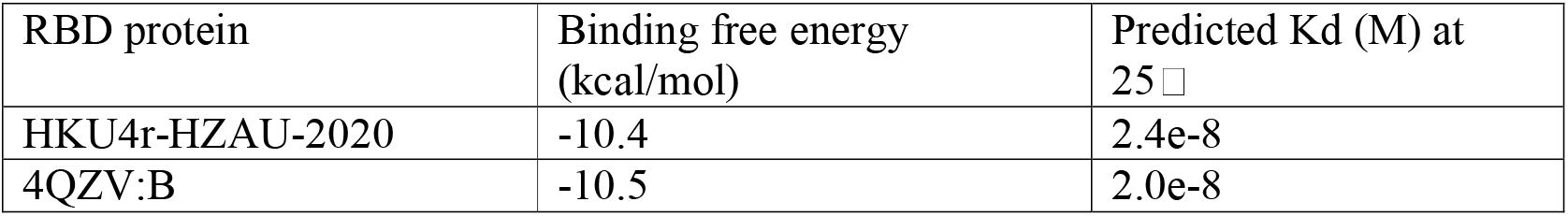
PRODIGY binding energy and predicted binding affinity of HKU4r-HZAU-2020 spike protein RBD and hDPP4 as compared to the RBD of HKU4 (PDB ID 4QZV).

We then compared seven HKU4-related and MERS-CoV sequences to HKU4r-HZAU-2020 in the key region of the RBD where interaction with human DPP4 occurs (Fig. 7). Six of the thirteen key hDPP4 binding residues have an exact match to MERS-CoV residues, while eleven of the thirteen key residues match those found in PDB:4QZV_B.

**Fig. 7.**
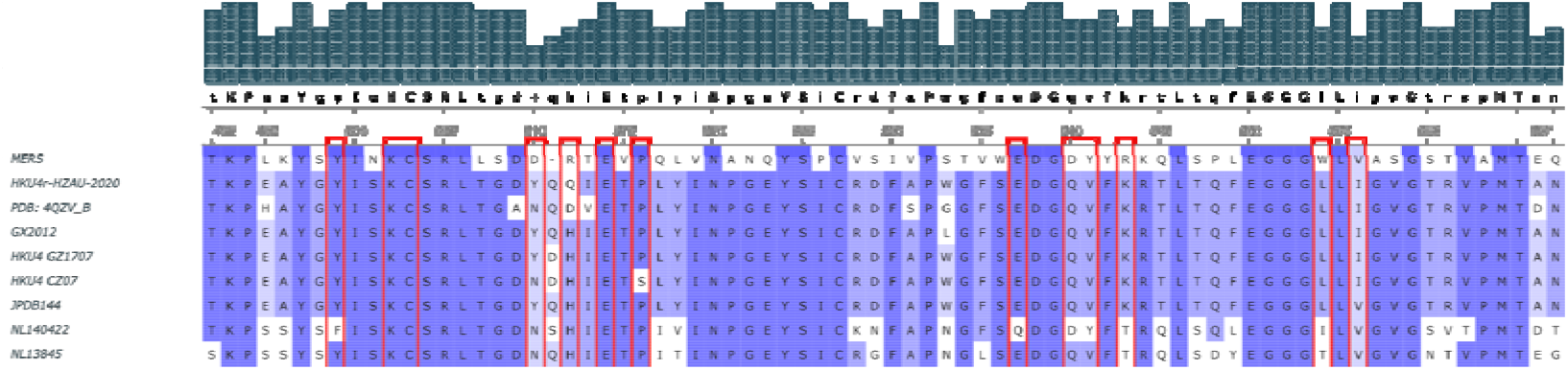
RBD subsequence alignment. Alignment of the external subdomain of the receptor binding domain for selected merbecoviruses. Key residue positions found to have direct interaction with human DPP4 (Luo et al., 2018) are marked with red bounding boxes. Plotted using UGENE. For protein accession numbers see Supp. Info. 1.7.

### MERS-CoV spike gene

From *de novo* assembly of the four pooled SRA datasets containing the novel HKU4-related CoV, we recovered two contigs of lengths 2,486 nt and 1,649 nt with 99.68% and 99.62% identity to the MERS-CoV reference strain HCoV-EMC/2012 (NC_019843.3), respectively (Supp. Info. 3.1). Coverage of the MERS S gene sequence by reads is 99.09%, however, the average read depth is low at 6.13 reads. A consensus C21695T SNV (read depth of 12) with respect to HCoV-EMC/2012 was found (Supp. Fig. 18). The only unmapped region of the MERS-CoV S gene is a 33 nt section located between positions 23,908 and 23,940 which is within the S2 subunit, upstream of the fusion peptide. Only four reads cover the 3’ most section of the spike sequence. Notably, all four reads have a 100% identity match to HKU4r-HZAU-2020 directly downstream of the MERS spike sequence (Supp. Figs. 18-19, Supp. Data). A 14 nt sequence at the 3’ end of the MERS S gene is identical to the equivalent position in the HKU4r-HZAU-2020 S gene, with the exception of C25514T (MERS-CoV reference) in HKU4r-HZAU-2020 (Supp. Fig. 20), as such the exact position of the join is uncertain.

At the 5’ end of the MERS-CoV S gene, three 150 nt reads with a 100% identity match to HCoV-EMC/2012 at their 3’ ends (44/44 nt, 78/78 nt and 125/125 nt), exhibit a 100% identity match to HKU4r-HZAU-20 at their 5’ ends (106/106nt, 72/72nt and 25/25 nt, respectively) (Supp. Fig. 21, Supp. Data). Two other reads aligning across the 5’ end of the MERS-CoV S gene exhibit only a single nucleotide variation (SNV) relative to HKU4r-HZAU-20 in a 25 nt section at their 5’ ends match, while at their 3’ ends a 36/36 nt match to the reverse complement of a 36 nt sequence at the 5’ end of the HCoV-EMC/2012 S gene is found (Supp. Fig. 21). We infer the 36 nt sequences at their 5’ ends likely represent ligation or reverse transcription step artifacts.

We infer that Golden Gate cloning was likely used for assembly of the HKU4r-HZAU-2020+S(MERS) chimera, since we did not identify any 1-8 cutter enzyme sequences in the MERS spike gene or HKU4r-HZAU-2020 in positions that could have been utilized for *in vitro* ligation. Additionally, we did not recover any synthetic vector sequences attached to the ends of the MERS spike sequences and no MERS-CoV sequences outside of the spike gene were found.

### Restriction enzymes and synthetic engineering detection

In an attempt to find an obvious signature of genetic engineering in the sequence of the HKU4r-HZAU-2020 genome, we performed a restriction enzyme mapping of the sequence using SnapGene using the set of all type II and type IIS restriction endonucleases with unique sites. Since BsmBI and BsaI are often used for the construction of coronavirus reverse genetic systems (Xie et al., 2020), we analyzed the number of these sites, as well as the number of unique restriction sites in the genome (Fig. 8). For comparison, we also obtained and performed similar restriction site mapping of two related coronaviruses: BtTp-BetaCoV/GX2012 (KJ473822.1) (Fig. 9) and HKU4 CZ07 (MH002338.1) (Fig. 10).

**Fig. 8.**
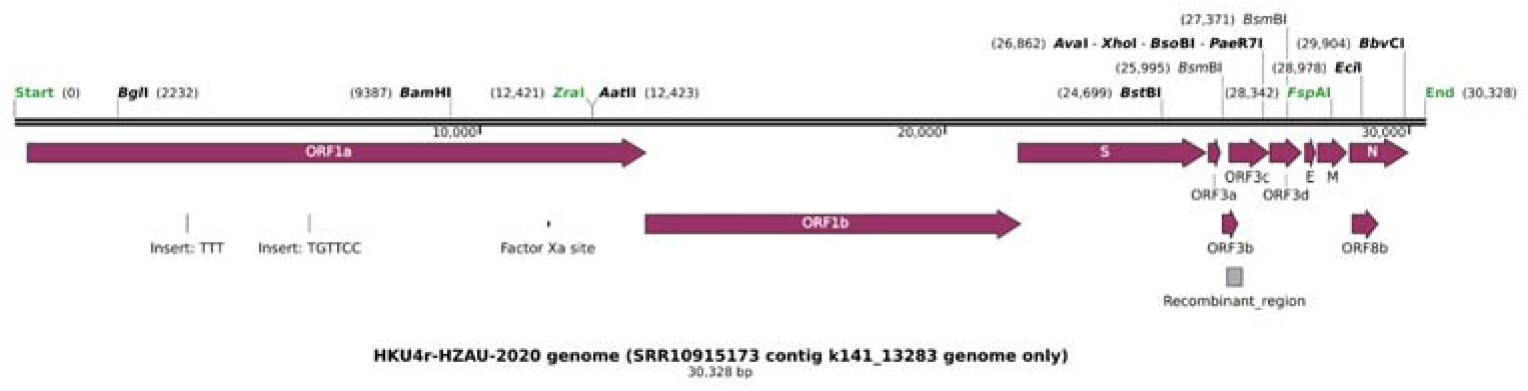
Annotation and restriction mapping of the HKU4r-HZAU-2020 genome.

**Fig. 9.**
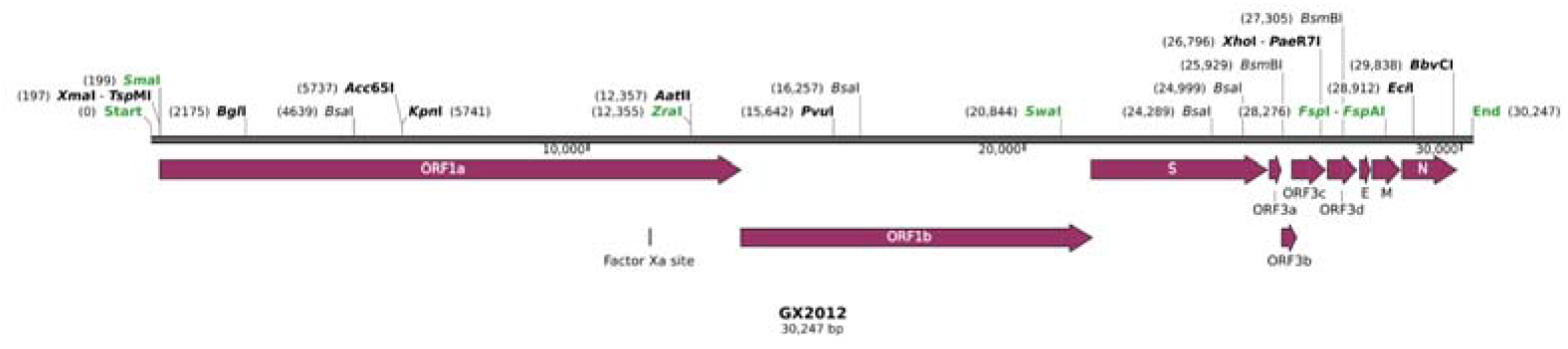
Annotation and restriction mapping of BtTp-BetaCoV/GX2012 for comparison with the HKU4r-HZAU-2020 genome.

**Fig. 10.**
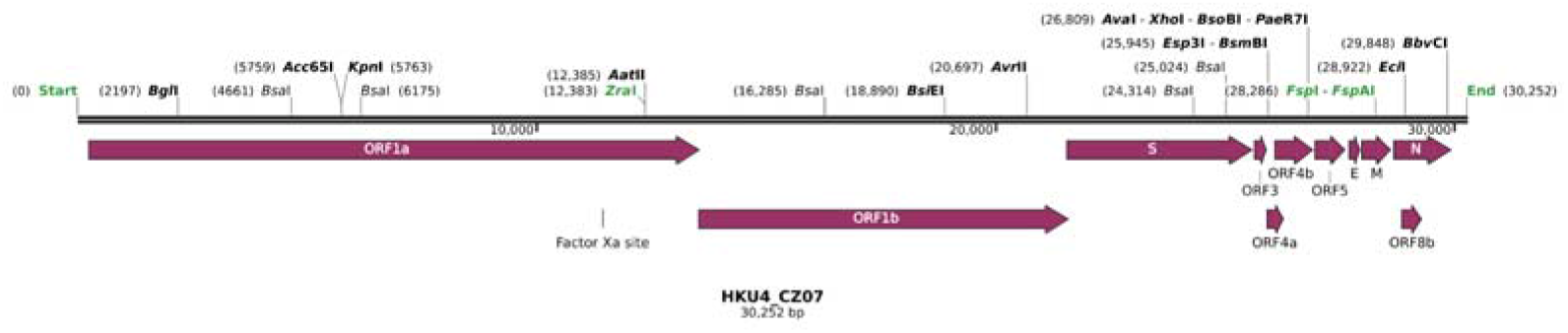
Annotation and restriction mapping of HKU4 CZ07 for comparison with the HKU4r-HZAU-2020 genome.

Several restriction sites were found to be conserved, including unique BglI, AatlI, XhoI, EciI, and BvbCI sites between two closely related HKU4-related CoV sequences and the HKU4r-HZAU-2020 genome (Fig. 12-14). However BsaI, BsmBI, and BamHI restriction sites across these sequences were not conserved. In the HKU4r-HZAU-2020 genome, there are only two BsmBI sites and a complete lack of BsaI sites. This appears anomalous, as all of the 14 most closely related HKU4-related CoVs, and MERS HCoV-EMC/2022 contain a minimum of five combined BsaI and BsmBI sites (Supp. Fig. 21, Supp. Table 1).

In an attempt to identify a potential previously known reverse genetics system for HKU4-related CoVs, we undertook a keyword search using “HKU4” and “reverse genetics” or “infectious clone” using PubMed and Google Scholar. However, we found no scientific publications which disclose such a reverse genetic system, or have demonstrated the construction of an infectious clone for HKU4-related coronaviruses. Consequently, HKU4r-HZAU-2020 represents a unique example of a BAC based full length HKU4-related CoV cDNA clone.

### Virus sequence contamination

Fastp filtered reads in each SRA dataset in BioProject PRJNA602160 were aligned to a set of viruses identified as present in the datasets as described in Methods. We identified six viral genomes and a synthetic sequence each with greater than 10% coverage in one or more SRA datasets in BioProject PRJNA602160 (Fig. 11; Supp. Fig. 22). As discussed above, HKU4r-CoV-HZAU and HCoV-EMC/2012 were only identified in four SRA datasets. These were the only RNA-sequencing datasets in PRJNA602160 using cDNA selection and a unicellular zygote source (Supp. Info. 1.1).

**Fig. 11.**
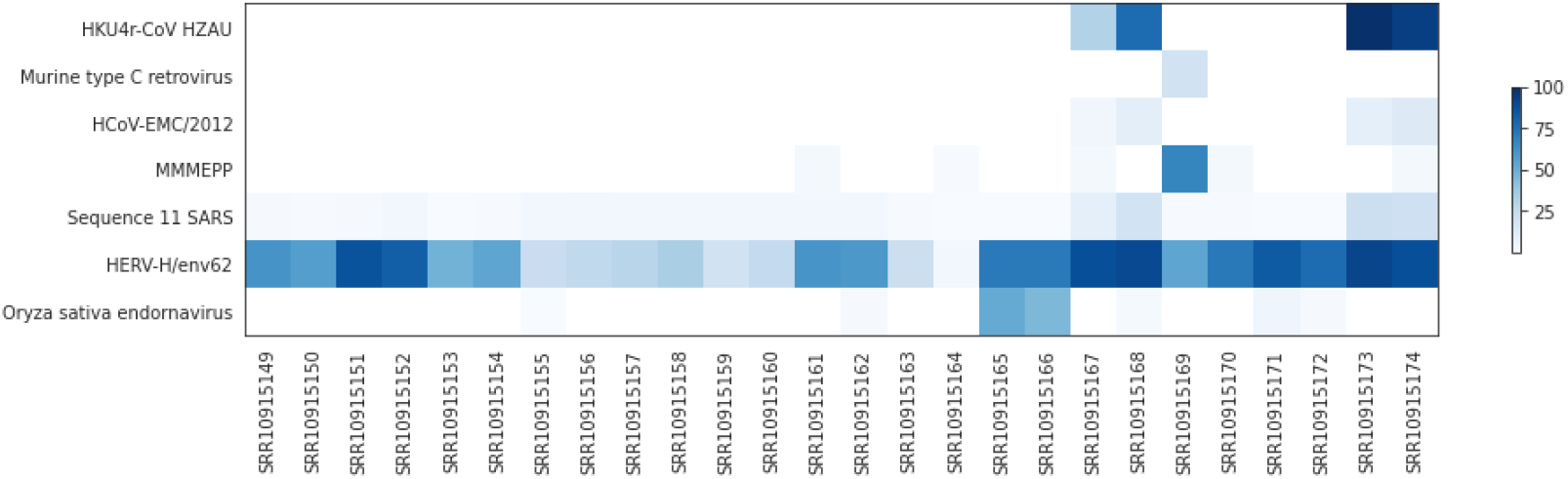
Percentage genome coverage for six viral genomes and one SARS BAC for each SRA dataset in PRJNA602160. A minimum cutoff of 10% coverage in any SRA was applied.

Human endogenous retrovirus H HERV-H/env62 sequences were identified in all SRA datasets. Murine hosted viruses were found in six SRA datasets but not obviously correlated with the presence of HKU4r-HZAU-2020.

The bacterial content in the four RNA-sequencing datasets containing HKU4r-HZAU-2020 sequences identified using NCBI STAT (Katz et al., 2021) analysis averaged 58.14%, with a range of 40.98% to 83.41% (Supp. Info. 1.2). Whereas the average bacterial content for the six RNA-sequencing datasets in BioProject PRJNA602160 which did not contain HKU4r-HZAU-2020 sequences averaged 1.65%, and contained a range of between 0.14% and 4.03%. We calculated a very strong spearman correlation coefficient of 0.81 between bacteria percentage and percentage of reads mapping to the HKU4r-HZAU-2020 viral genome in RNA-sequencing datasets in BioProject PRJNA602160 (Supp. Code).

### Negative control

We identified that RNA-Sequencing BioProject PRJNA602115, a zebrafish miR-462-731 regulation study (Huang et al., 2020) was submitted by the HZAU to NCBI on the same day as BioProject PRJNA602160. As a control dataset, we ran the same workflow we used for identifying and aligning HKU4r-HAZU-2020 on the three SRA datasets in BioProject PRJNA602115. Only a small number of viruses were identified using fastv, all with low coverage (Supp. Info. 4.1). We used minimap2 to align the three SRA datasets to the same set of HKU4-related CoVs, MERS-CoV, and miscellaneous viruses used for PRJNA602160 contamination identification, as well as the three viruses identified using fastv with more than 3% coverage (Supp. Info. 4.2). Only Sugarcane mosaic virus (NC_003398.1) and the Harvey murine sarcoma virus p21 v-has protein gene (NC_038668.1) were found to have more than 10% coverage. No trace of HKU4-related CoVs or MERS-CoV was found (Supp. Info. 4.2). Additionally, we aligned the three SRA datasets to all mitochondrial sequences on NCBI. As expected, the *Danio rerio* mitochondrial gene was mapped with near complete coverage in all datasets. We additionally identified *Ochotona curzoniae* and *Rattus norvegicus* genomic sequence contamination in the datasets (Supp. Info. 4.3).

## Discussion

SARS-related and MERS-related coronaviruses (CoVs) with zoonosis potential have been a subject of intense research since the SARS-CoV and MERS-CoV spillovers to humans in 2002 and 2012, respectively. In addition to MERS-related CoVs, the two other groups belonging to the merbecovirus subgenus of betacoronaviruses are HKU4-related and HKU5-related CoVs. Here we have identified a full-length HKU4-related CoV cDNA clone contaminating an *Oryza sativa japonica* agricultural sequencing BioProject. The average read depth of 66 for the clone in pooled datasets is good, and we have high confidence in both the presence of the clone as well as its sequence.

HKU4-related CoVs are only known to be hosted by *Tylonycteris* bats (Vespertilionidae family, Vespertilioninae subfamily), a rare insectivorous genus only known to live in bamboo forests, and in China only found in the Guangdong, Guangxi, Yunnan, Guizhou, Hong Kong, and Macau regions of southern China (Fan et al., 2019). HKU4r-HZAU-2020, the novel HKU4-related CoV identified here, phylogenetically groups with *Tylonycteris pachypus* isolate BtTp-BetaCoV/GX2012 sampled from Guangxi province and *Tylonycteris* bat HKU4 isolates CZ07 and CZ01 over the full genome, S and RdRp genes. HKU4 isolates CZ07 and CZ01 were sampled from southern China by WIV-affiliated researchers in 2012 (Supp. Info. 1.4). BtTp-BetaCoV/GX2012, which exhibits highest identity over the full genome, was sampled by Wu et al. (2016) in the Guangxi province in 2012 (Supp. Info. 1.4). Tylonycteris bat coronavirus HKU4 isolate 152762, with closest partial RdRp sequence grouping with HKU4r-HZAU-2020, was sampled between 2010 and 2015 by Latine et al. (2020) as part of research co-authored by WIV-affiliated researchers (Supp. Info. 1.8). Consequently, we infer that the HKU4r-HZAU-2020 genome is either a *Tylonycteris* sp. hosted CoV sampled by researchers from southern China, or is a consensus genome generated from *Tylonycteris* spp. hosted CoVs.

Both the novel HKU4-related CoV genome and its corresponding reverse genetics system were previously undocumented. Moreover, no reverse genetics systems were previously published for any HKU4-related CoVs. It is interesting to note that a 5-year grant awarded to EcoHealth Alliance (EHA) by the National Institute of Health (NIH) involved chimeric MERS-related CoV construction at the WIV (Daszak, 2021). Grant award year 5 (2018-06-01 – 2019-05-31) included generating full-length infectious clones of MERS-CoV with the RBD replaced by those from various HKU4-related CoVs and assessing the infectivity of novel chimeras in human cells. Two full MERS-related CoV genomes from Guangdong province, BtCoV/li/GD/2013–845 and BtCoV/li/GD/2014–422, were published by Luo et al. (2018) and discussed in Daszak (2021), while notably 24 HKU4-related CoV sequences were not published.

We find that instead of using a published MERS-CoV backbone and inserting novel HKU4-related CoV fragments (Daszak, 2021), a completely undocumented HKU4-related CoV clone was apparently being researched in Wuhan. The discovery of a near complete MERS spike sequence in the same BioProject as HKU4r-HZAU-2020 is concerning. Although we identified only a low number of reads mapping across the 5’ and 3’ ends of the MERS spike gene sequence, these reads clearly indicate that a MERS-CoV S gene sequence had been inserted into the HKU4r-HZAU-2020 backbone. The upstream section of reads mapping across the 5’ end of the MERS S gene, had a 100% match part of the ORF1b gene of HKU4r-HZAU-2020 genome directly upstream of its spike gene. Similarly, the downstream section of reads mapping across the 3’ end of the MERS S gene, had a 100% match to a non-coding region of the HKU4r-HZAU-2020 genome directly downstream of its spike gene. The probability that such a configuration could be artificial we believe is extremely remote. As such we infer it highly probable that a second HKU4-related CoV clone ‘HKU4r-HZAU-2020+S(MERS)’, with a replacement of the spike sequence by a MERS-CoV spike is present in the BioProject, albeit at lower abundance than HKU4r-HZAU-2020.

The lack of BsaI sites and the low combined count of BsaI and BsmBI sites are both anomalous in HKU4r-HZAU-2020 relative to the 14 other HKU4-related CoVs (Supp. Table 1). We infer it is likely that “No See’m” cloning (Yount et al., 2022) or Golden Gate assembly (Engler et al., 2008) was used to assemble the HKU4r-HZAU-2020 genome fragments, whereby fragments can be ligated in one step for a seamless assembly (Engler and Marillonnet, 2014).

Merbecoviruses were actively researched in Wuhan, predominantly by the WIV with documented plasmid construction, genetic engineering and viral entry studies (Yang et al., 2015; Sun et al., 2017; Luo et al., 2018, Xia et al., 2019; Zhang et al., 2019; Daszak 2021). Indeed we note that the synthetic vector backbone of pBAC-SARS-CoV, which best matches the synthetic sequences present at the 3’ end of HKU4r-HZAU-2020 and shows 100% sequence matches of 956 nt and 587 nt lengths at the 5’ end, was used previously by researchers at the WIV (Wang et al. 2008). However, merbecovirus research has also been conducted by HZAU researchers (Chen et al., 2017, Yuan et al., 2017) and Wuhan University (Wu et al., 2015; Han et al., 2017). As the WIV outsourced CoV sequencing to Wuhan sequencing centers (Cohen, 2020), the originating laboratory for HKU4r-HZAU-2020 is uncertain.

Cross-contamination of samples is a well-documented challenge in biological sequencing laboratories, as noted by several studies including those by Lusk (2014) and Selitsky et al. (2020). In particular, Cantalupo and Pipas (2019) have highlighted several key stages in the sequencing process where contamination can occur, including sample collection and shipping, where “passenger” viruses can cross-contaminate, as well as during virus purification, nucleic acid isolation, amplification, and library preparation, where there is a risk of reagent contamination and amplification bias. Additionally, contamination may also occur due to human error, such as improper use of protective equipment or failure to follow established protocols, or due to laboratory infrastructure issues, such as poor ventilation systems or inadequate physical separation of samples. Contamination can also occur during sequencing itself, through machine contamination, index hopping, and sequencing errors.

HKU4r-HZAU-2020 is manifestly a full-length cDNA clone engineered as a BAC, and as such would be unrelated to RNA contamination. We found that all RNA-sequencing datasets in rice BioProject PRJNA602160 without merbecovirus sequences had a maximum of 4% bacteria, while the minimum quantity of bacteria contaminating the four SRA datasets containing merbecovirus reads was 41%. This finding is strongly indicative of upstream contamination, prior to sequencing. The RNA-sequencing of samples in BioProject PRJNA602160 involved a specialized library format and protocol. While we cannot be certain, given this specific library format and protocol, we infer it unlikely that another library was prepared using the same format and pooled in the same run as samples in BioProject PRJNA602160. RNA-seq of the rice dataset was conducted with SMART-seq V4 Ultra low input RNA kit, sheared with AMPure beads, and the resulting 200-400 bp cDNA prepared with a TruSeq CHIP DNA kit. The RNA was isolated directly from zygotes collected with a micromanipulator system using FDA staining, and the resulting single cells were constructed individually with one library for each cell. The micromanipulator and staining prior to direct RNA extraction and input into a low-input cDNA kit, followed by a unique library construction, implies that the protocol was conducted in a specialized agricultural laboratory with no external laboratory work for the isolation through sequencing stages. As such, while the source of the HKU4-HZAU-2020 clone and MERS-CoV sequences could be due to index hopping from a multiplexed run, we infer it to be unlikely.

We propose that the most likely scenario of contamination of SRR10915167-8 and SRR10915173-4 is HKU4r-HZAU-2020 plasmids contaminating the samples prior to sequencing. Similarly, we infer that contamination by MERS spike gene sequences likely occurred via HKU4r-HZAU-2020+S(MERS) plasmids also contaminating the samples prior to sequencing. Contamination by plasmids prior to sequencing is supported by the finding of a very high spearman correlation coefficient between bacterial content and the number of reads mapping to the HKU4r-HZAU-2020 genome in RNA-sequencing datasets in BioProject PRJNA602160 (Supp. Code).

The BioProject for the *Oriza sativa japonica* sequencing datasets was registered at GenBank on 2020-01-19, but to this date the novel HKU4-related CoV has not been published by the HZAU or WIV. We note that *Oryza sativa* cultivar:japonica (rice) sequencing BioProject PRJNA601977 was registered on NCBI on 2020-01-17 by the WIV, two days before the registration of the *Oryza sativa japonica* BioProject PRJNA602160 by the HZAU (containing the novel HKU4-related CoV clone), indicating the two projects may be related, however this observation cannot be confirmed as no data has been published for PRJNA601977 by the WIV.

The potential for bat-hosted coronaviruses to spill over to humans and spark epidemics has been widely documented and is a subject of extensive research (Hu et al., 2017; Yu et al., 2019; Epstein et al., 2020; Lau et al., 2021). Our finding that HKU4r-HZAU-2020 likely binds to hDPP4 adds to previously documented research that two HKU4-related CoVs are able to bind to hDPP4. HKU4 strain B04f (EF065505.1) from Guangdong province and HKU4 strain SM3A were each able to bind hDPP4 (Yang et al., 2014; Wang et al., 2014; Lu et al., 2020, Lau et al., 2021). However, HKU-related-CoVs including HKU4r-HZAU-2020 do not possess a human proprotein convertase cleavage motif at the S1/S2 boundary. As such we infer that HKU4r-HZAU-2020 may be incapable of mediating sustained human cell entry without the evolution of, or introduction of a furin cleavage motif (Yang et al., 2015).

A topic which receives far less attention than zoonotic spillover is the emerging threat of accidental human infection during laboratory research of wild type viruses, or research involving enhanced potential pandemic pathogens (Butler, 2015; Klotz, 2019; Shinomiya et al., 2022). Although the exact nature of the research related to the HKU4r-HZAU-2020 clone is unknown, there are two experimental purposes in generating infectious clones of coronaviruses: to study a wildtype virus for which no ‘live’ isolate exists, but for which a genome sequence is available, or to manipulate the genome of a coronavirus in order to study changes in infectivity, tropism (including host or tissue specificity), or pathogenicity.

Regarding the potential connection of our findings to uncovering the origin of SARS-CoV-2, one of the main arguments that has been adduced for the hypothesis that SARS-CoV-2 or any potential precursors could not have been manipulated in a laboratory, is the absence of a published backbone closely related to SARS-CoV-2. Liu et al. (2020) claimed that because SARS-CoV-2 is too differentiated from SL-SHC014-MA15, a published mouse adapted chimeric CoV generated using a SARS backbone (Menachery et al., 2015), that SARS-CoV-2 could not have been engineered. In addition, Andersen et al. (2020) wrote: “*if genetic manipulation had been performed, one of the several reverse-genetic systems available for betacoronaviruses would probably have been used*”. Both arguments are clearly undermined by our findings. The notion that all reverse-genetic cloning systems in contemporaneous use for research had been made known to the scientific community was already tenuous, now it should be abandoned completely in light of these discoveries. There are no known publications documenting a reverse genetic system for a HKU4-related CoV strain.

Bat CoVs HKU4 and HKU5 are related to MERS CoVs (Wang et al., 2014). As MERS-CoV is a BSL-3 level pathogen (Teramichi et al., 2019) with approximately 36% mortality (World Health Organization, 2022), MERS-related CoV research should be a highly regulated process, conducted within high-containment biosafety laboratories. The finding of a HKU4-related full length CoV clone with *in silico* predicted hDPP4 binding ability (HKU4r-HZAU-2020) and HKU4r-HZAU-2020+S(MERS), a likely clone with a substituted MERS-CoV spike in agricultural research datasets is troubling. Agricultural sequencing facilities do not have the same high biosafety standards or protocols as BSL-3 laboratories (Klein, 2012; Heckert and Kozlovac, 2014), and as such the potential for accidental human contact with infectious agents is significantly higher. To reduce the risk of another pandemic, we strongly recommend strict international regulation and oversight of enhanced potential pandemic pathogen research.

## Conclusion

In this work, we have reported the unexpected discovery of a HKU4-related coronavirus (CoV) clone in an agricultural rice sequencing data from the Huazhong Agricultural University in Wuhan. The novel HKU4r-HZAU-2020 genome exhibits a 98.38% identity to the closest published HKU4-related CoV, BtTp-BetaCoV/GX2012. The regions of most significant difference to the most closely related HKU4-related CoVs are in the spike, ORF3a and ORF3b coding regions. We modeled the interface of the RBD of the novel HKU4-related CoV to human dipeptidyl peptidase 4 (hDPP4) and found that HKU4r-HZAU-2020 likely binds to human cells. This represents the third known HKU4-related CoV with hDPP4 binding potential.

The HKU4r-HZAU-2020 genome was found to have been inserted into a bacterial artificial chromosome (BAC). T7 promoter and cytomegalovirus promoter sequences were identified upstream of the genome, likely to facilitate *in vitro* transcription of full length RNA copies of the genome. Downstream of the 3’ end of the poly(A) tail, a hepatitis delta virus ribozyme and bovine growth hormone termination and polyadenylation sequence were found, which would allow truncation of the 3’ end of the transcribed viral genome. This constitutes the first known reverse genetics system used for HKU4-related CoV research.

We further identified the presence of a near complete MERS-CoV spike sequence which had been inserted into the HKU4r-HZAU-2020 backbone forming a second clone in the rice RNA-sequencing datasets. As the MERS-CoV RBD binds more efficiently to hDPP4 than known HKU4r-CoVs, and as the MERS-CoV S protein has the demonstrated capability of utilizing human cell proteases for mediating cell entry, the HKU4r-HZAU-2020+S(MERS) chimera appears to constitute enhanced potential pandemic pathogen (gain-of-function) research.

We infer the most likely source of contamination of the rice RNA-sequencing datasets was by bacterial plasmids upstream of the sequencing step. Our findings highlight the potential biosafety risks of enhanced potential pandemic pathogen research, whereby human error, even during routine tasks such as sequencing, could potentially lead to inadvertent laboratory-acquired infection with dangerous human pathogens.

Finally, this work serves as a cautionary story to show that the absence of a documented related reverse genetic system cannot be relied upon as evidence to disprove laboratory involvement of a novel viral pathogen.

## Materials & Methods

### Datasets

Analysis of each SRA experiment in *Oryza sativa japonica* group sequencing project accession PRJNA602160 in the NCBI SRA database (Agarwala et al., 2018) was conducted using the NCBI SRA Taxonomy Analysis Tool (STAT), a k-mer-based taxonomic classification tool (Katz et al., 2021). BtTp-BetaCoV/GX2012 (KJ473822.1) was identified as a genome with significant matches in SRA datasets: SRR10915168, SRR10915173, and SRR10915174. SRR10915167 exhibited matches to *Tylonycteris* bat coronavirus HKU4 (NC_009019.1) and MERS-CoV (NC_019843.3).

SRA format datasets from NCBI were extracted as single fastq files using sratoolkit version 3.0.0 (https://trace.ncbi.nlm.nih.gov/Traces/sra/sra.cgi?view=software).

Fastv (Chen et al., 2020) was run for each SRA against the Opengene vial genome kmer collection ‘microbial.kc.fasta.gz’ (https://github.com/OpenGene/UniqueKMER). Four datasets that contained HKU4-related CoV sequences and three datasets that contained MERS CoV sequences at greater than 5% genome coverage were identified (Supp. Info. 1.8).

Each SRA dataset in PRJNA602160 was filtered using fastp (Chen et al., 2018) with default settings, and filtered datasets were used for all subsequent analysis.

### Alignment and Assembly

Read alignments were conducted using Minimap2 version 2.24 (Li et al., 2018) with the following parameters unless indicated “-MD -c -eqx -x sr --sam-hit-only -- secondary=no -t 32”.

Alignments of pooled reads from SRR10915167, SRR10915168, SRR10915173, and SRR10915174 to the MERS reference sequence HCoV-EMC/2012 (NC_019843.3) was made using bowtie2 version 2.4.2 (Langmead and Salzberg 2012) using the ‘--local’ setting.

MEGAHIT v1.2.9 (Li et al., 2015) with default settings was used for de novo assembly of each SRA dataset. To confirm the MEGAHIT assembly, we also undertook *de novo* assembly of SRR10915167-8 and SRR10915173-4 using coronaSPAdes v3.15.2 (Meleshko et al., 2021) using default settings, and SPAdes v3.15.2 (Prjibelski et al., 2020) using the ‘--careful’ parameter. Additionally SRR10915167-8 and SRR10915173-4 were pooled and also assembled with coronaSPAdes and SPAdes with settings as per above.

*De novo* assembled contigs were aligned using minimap2 to a concatenated virus database consisting of the NCBI viral database downloaded on 2021-11-28 and all CoVs on NCBI downloaded on 2020-03-30 (https://github.com/ababaian/serratus/wiki/Working-Data-Dir). Contigs from SRR10915173 were aligned using minimap2 to the NCBI nt database downloaded on 2021-11-27. A set of viruses with greater than 5% coverage after fastv read analysis and contig alignments to NCBI databases was then generated. Each SRA dataset in PRJNA602160 was aligned using minimap2 to this selected set of viruses.

The 38,583 nt contig from MEGAHIT *de novo* assembly of SRR10915173 ‘k141_13282’ was trimmed to remove the first 149 nt. Reads from SRR10915167, SRR10915168, SRR10915173 and SRR10915174 were aligned to the resultant 38,434 nt trimmed contig ‘k141_13282_del_149’ using minimap2. Additionally, pooled reads from the four SRA datasets above were aligned to the HKU4r-HZAU-2020 clone (i.e. contig ‘k141_13282_del_149’) using bowtie2 and bwa-mem2 (Vasimuddin et al., 2019) using default settings.

Reads in all SRAs were aligned to the entire NCBI mitochondrial database downloaded on 2022-05-05, using minimap2.

Mapping statistics and coverages for all alignments were calculated using samtools version 1.15.1 (Li et al. 2009) and bamdst version 1.0.9 (Bamdst, 2021).

### Phylogenetic and recombination analyses

For all phylogenetic analyses, the following workflow was used. The novel CoV genome, and genome sub-regions were analyzed using blastn (Johnson et al., 2008) to identify the 100 closest genomes. These were downloaded, and aligned using MAFFT v7.490 (Katoh and Standley 2013) using the ‘auto’ parameter with default settings. A PhyML (Guindon et al., 2010) maximum likelihood tree was generated with smart model selection (SMS) (Lefort et al., 2017) using default settings. Tree branches distant from the query genome were pruned. A maximum-likelihood tree using the pruned genome set was then generated using PhyML with SMS using default settings. The result was then plotted using MEGA11 (Tamura et al., 2021). Additionally for each pruned genome set MEGA11 model selection was used to identify the ML model with lowest bayesian information criterion (BIC). A maximum likelihood tree was generated in MEGA11 using this model with 1,000 bootstrap replicates unless otherwise indicated.

RDP4 (Martin et al., 2015) was used with with the following algorithms to test recombination regions in the HKU4r-HZAU-2020 genome: RDP method (Martin and Rybicki 2000), BOOTSCAN (Salminen et al., 1995), MAXCHI (Maynard Smith 1992), CHIMAERA (Posada and Crandall 2001), 3SEQ (Boni et al., 2007), GENECONV (Padidam et al. 1999), LARD (Holmes, Worobey, and Rambaut 1999), and SISCAN (Gibbs, Armstrong, and Gibbs 2000). The HKU4r-HZAU-2020 genome was tested against 13 closely related HKU4-related CoV genomes identified using blast against the nt database: BtTp-BetaCoV/GX2012; HKU4 isolate CZ07; HKU4 isolate CZ01; BtCoV/133/2005; HKU4 isolate SM3A; HKU4-4; HKU4-related isolate GZ131656; HKU4 isolate SZ140324; HKU4-2; HKU4-3; HKU4; Bat CoV isolate JPDB144 and Tr CoV isolate 162275. See Supp. Info. 1.4 for GenBank accession numbers.

### Sequence annotation and restriction site mapping

Gene and vector sequence annotation was conducted using ApE version 3.1.4 (Davis et al., 2022) to identify open reading frames and SnapGene version 6.2 (SnapGene®) for feature annotation. Restriction enzyme site annotation was conducted in SnapGene version 6.2.

### Modeling the Receptor Binding Domain

Structure of the RBD for the novel HKU4-related CoV was modeled using SWISS-MODEL web server (Waterhouse et al., 2018) and aligned to PDB id: 4QZV using PyMol Version 2.4 (PyMOL). Contacts were identified using default distance settings. The binding free energy of the complex was calculated using PRODIGY web server (Xue et al., 2016) and compared against that of the canonical complex PDB id: 4QZV.

### Nucleotide sequence accession number

The final HKU4r-HZAU-2020 clone sequence was submitted to GenBank as accession number OK560913.

## Acknowledgements

We thank Francisco A. de Ribera, Valentin Bruttel, Adrian Gibbs and Jonathan Latham, as well as an anonymous reviewer, for review and helpful comments.

## Data availability

Supplementary information and data are available on Zenodo:

doi: 10.5281/zenodo.7633114

link: https://zenodo.org/record/7633114

### Supplementary Information

Supp_Info_1.xlsx: BioProject PRJNA602160 SRA analysis.

Supp_Info_2.xlsx: Mitochondrial sequence analysis for all SRA datasets in BioProject

PRJNA602160.

Supp_Info_3.xlsx: MERS-CoV sequence analysis from SRA datasets in BioProject PRJNA602160.

Supp_Infor_4.xlsx: Analysis results for control datasets in PRJNA602115.

### Supplementary Data

Snapgene annotated image of HKU4r-HZAU-2020 clone:

SRR10915173_megahit_default_fastp_HKU4r-CoV_IC_k141_13283_del_149bp_Map.png

HKU4r-HZAU-2020 complete sequence in fasta format: HKU4r-HZAU_2020_SRR10915173_k141_13283_del_149.fa

HKU4r-HZAU-2020 annotated genome in GenBank format: SRR10915173_megahit_default_fastp_HKU4r-CoV_IC_k141_13283_del_149bp.gbk

HKU4r-HZAU-2020 genome only without synthetic vectors in fasta format: HKU4r-HZAU-2020_complete_genome_only.fa

HKU4r-HZAU-2020 annotated genome only in GenBank format: SRR10915173_megahit_default_fastp_HKU4r-CoV_IC_k141_13283_del_149bp_genome_only.gbk

Contigs with high identity to the MERS-CoV genome (NC_019843.3) assembled from pooled SRAs SRR10915167, SRR10915168, SRR10915173, SRR10915174 in fasta format: MEGAHIT_MERS_NC_19843.3_spike_contigs.fa.

Three-dimensional model of the RBD of the HKU4r-HZAU-2020 clone: HKU4r-HZAU-2020_RBD.pdb

Reads in SRR10915167-8 and SRR10915173-4 mapping to the HKU4r-HZAU-2020 clone using minimap2:

SRR10915167-68_73-74_k141_13283_rc_del_149_minimap2_samclip_25.bam

CoronaSPAdes default *de novo* assembled contigs aligned to the HKU4r-HZAU-2020 clone using minimap2: SRR10915173_coronaspades_default_k141_13283_rc_del_149_minimap2.bam

SPAdes careful *de novo* assembled contigs aligned to the HKU4r-HZAU-2020 clone using minimap2:

SRR10915173_spades_careful_k141_13283_rc_del_149_minimap2.bam

metaxa2 rRNA sequences de novo assembled using MEGAHIT: metaxa2_contigs.zip

Reads from pooled datasets SRR10915167-8 and SRR10915173-4 aligned to S gene section of MERS-CoV genome (NC_019843.3):

SRR10915167-68_73-74_NC_019843_3_S_gene_minimap2.sam

Reads mapping to MERS-CoV (NC_019843.3) S gene and crossing it’s 5’ and 3’ ends showing S gene insertion into HKU4r-HZAU-2020 backbone:

NC_19843.3_spike_gene_3prime_read_stats.txt

NC_19843.3_spike_gene_5prime_read_stats.txt

### Supplementary Code

Python code for image generation workflows is available at: https://github.com/bioscienceresearch/Discovery_of_Merbecovirus_clone

## Supplementary Figures

**Supp. Fig. 1.**
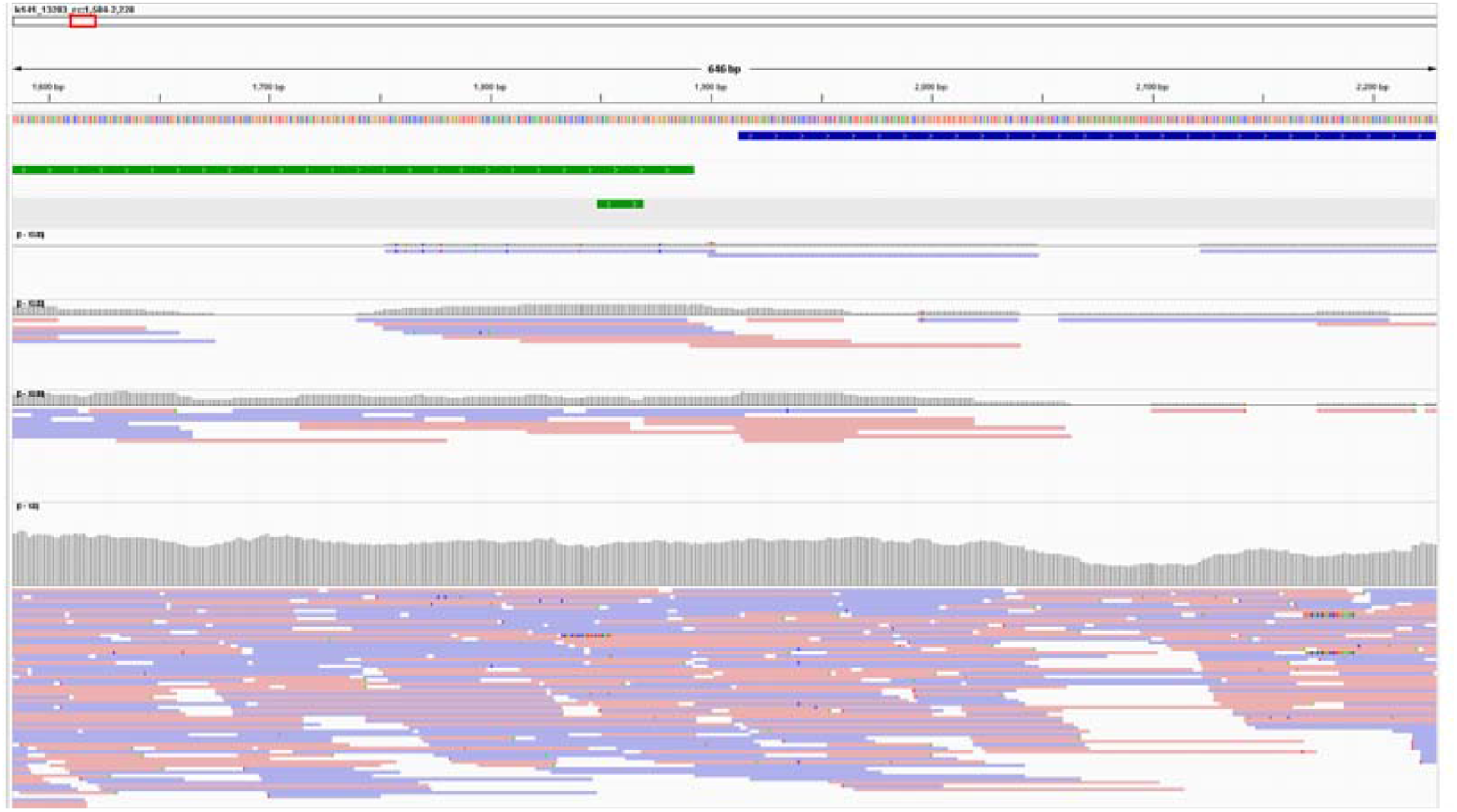
Read depth coverage at the 5’ end of the HKU4r-HZAU-2020 genome for each of the four SRAs containing HKU4r-HZAU-2020 sequences. From top to bottom: a) HKU4r-HZAU-2020 genome sequence (blue); b) CMV promoter sequence (green); c) CMV forward primer sequence; d-g) read depth (gray) and reads mapping to contig k141_13283 for SRR10915167, SRR10915168, SRR10915174 and SRR10915173. Read depths for SRR10915173 shown in 0-100 scale, for all other SRA datasets, read depth shown in 0-10 scale.

**Supp. Fig. 2.**
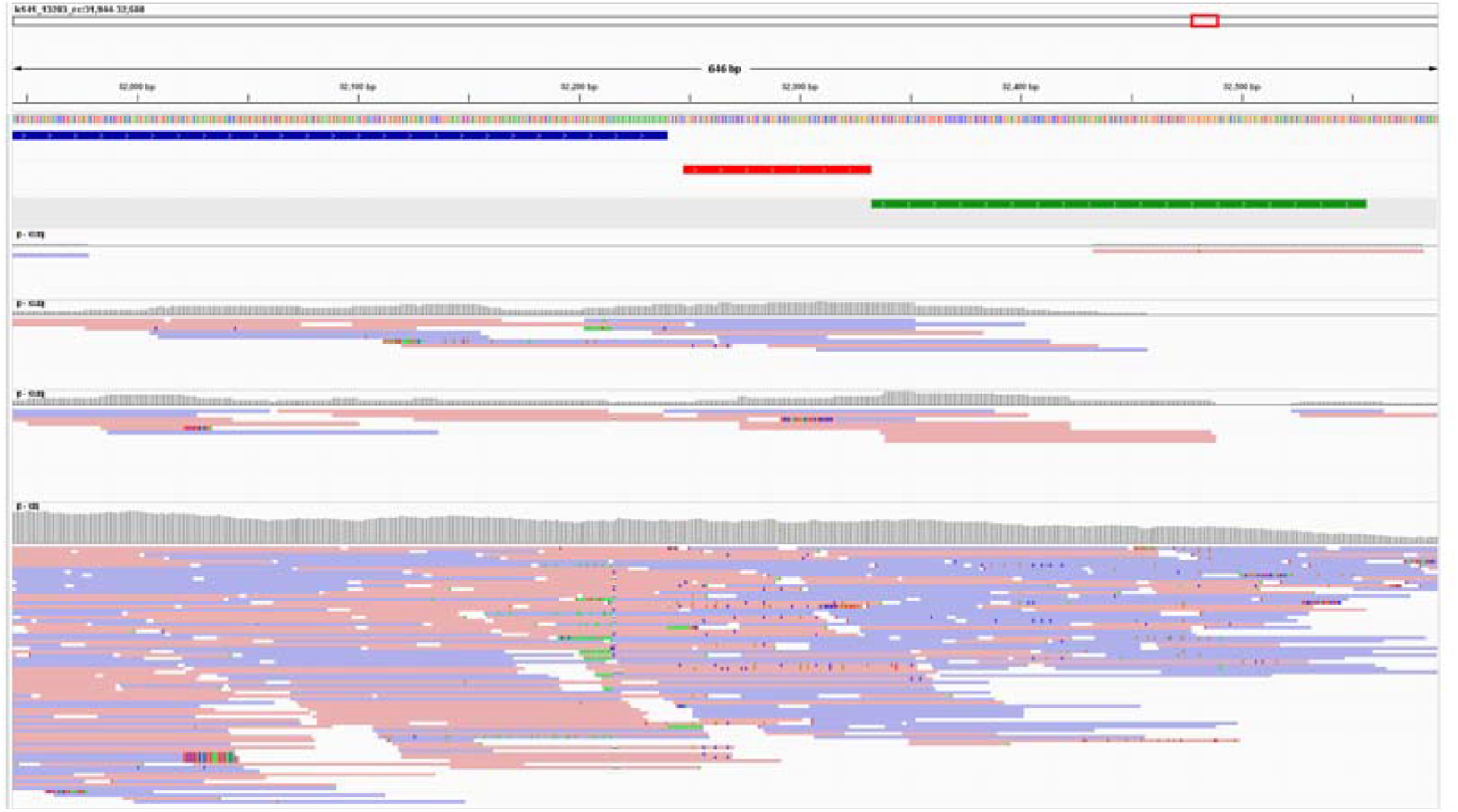
Read depth coverage at the 3’ end of the HKU4r-HZAU-2020 genome for each of the 4 SRAs containing HKU4r-HZAU-2020 sequences. From top to bottom: a) HKU4r-HZAU-2020 genome sequence (blue); b) HDV ribozyme terminator (red); c) bGH poly(A) signal (green); d-g) read depth (gray) and reads mapping to contig k141_13283 for SRR10915167, SRR10915168, SRR10915174 and SRR10915173. Read depths for SRR10915173 shown in 0-100 scale, for all other SRA datasets, read depth shown in 0-10 scale.

**Supp. Fig. 3.**
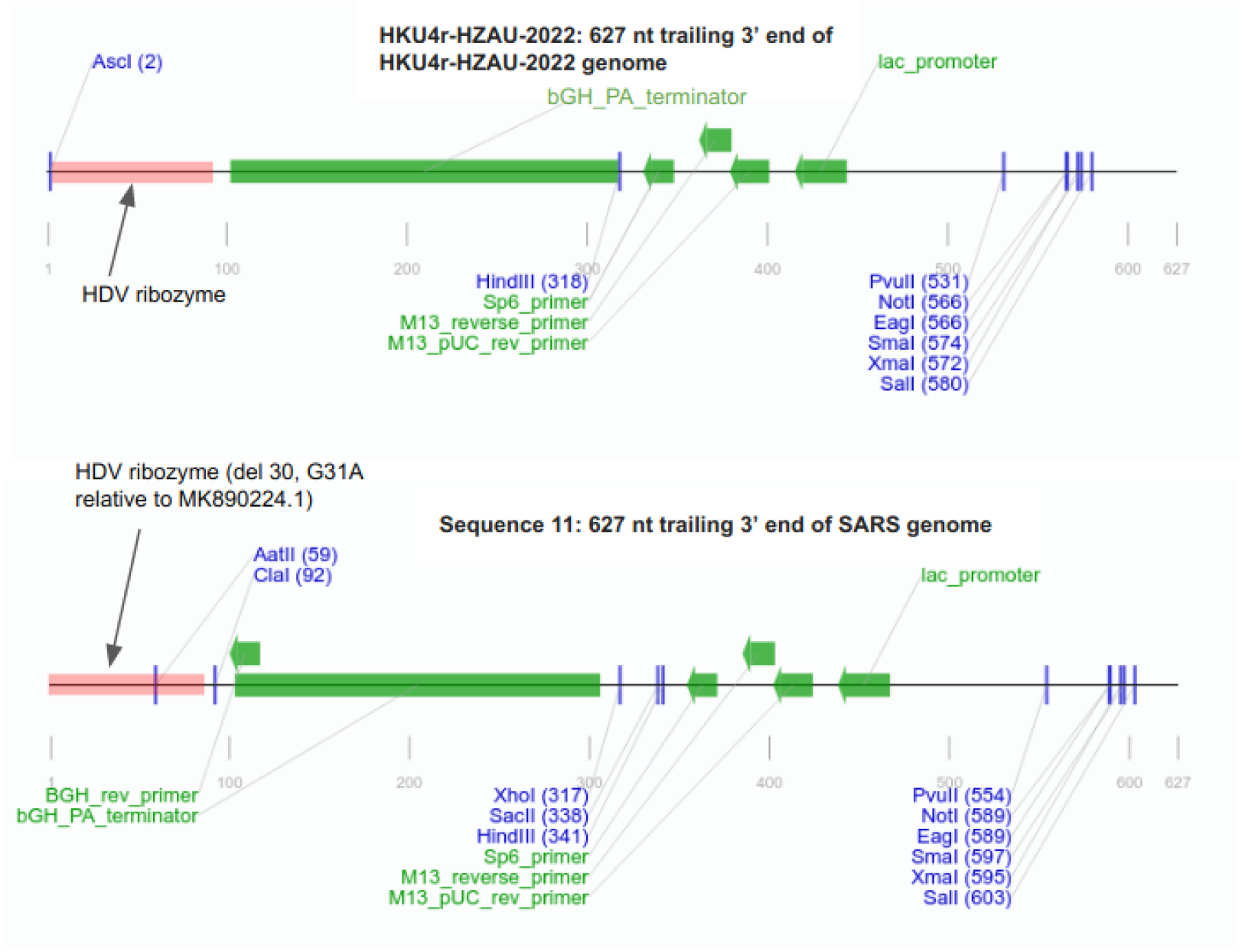
627 nt sequence trailing HKU4r-HZAU-2020 genome as compared with pBAC-SARS-CoV Sequence 11 from Patent WO2006236448 (accession CS480537.1).

**Supp. Fig. 4.**
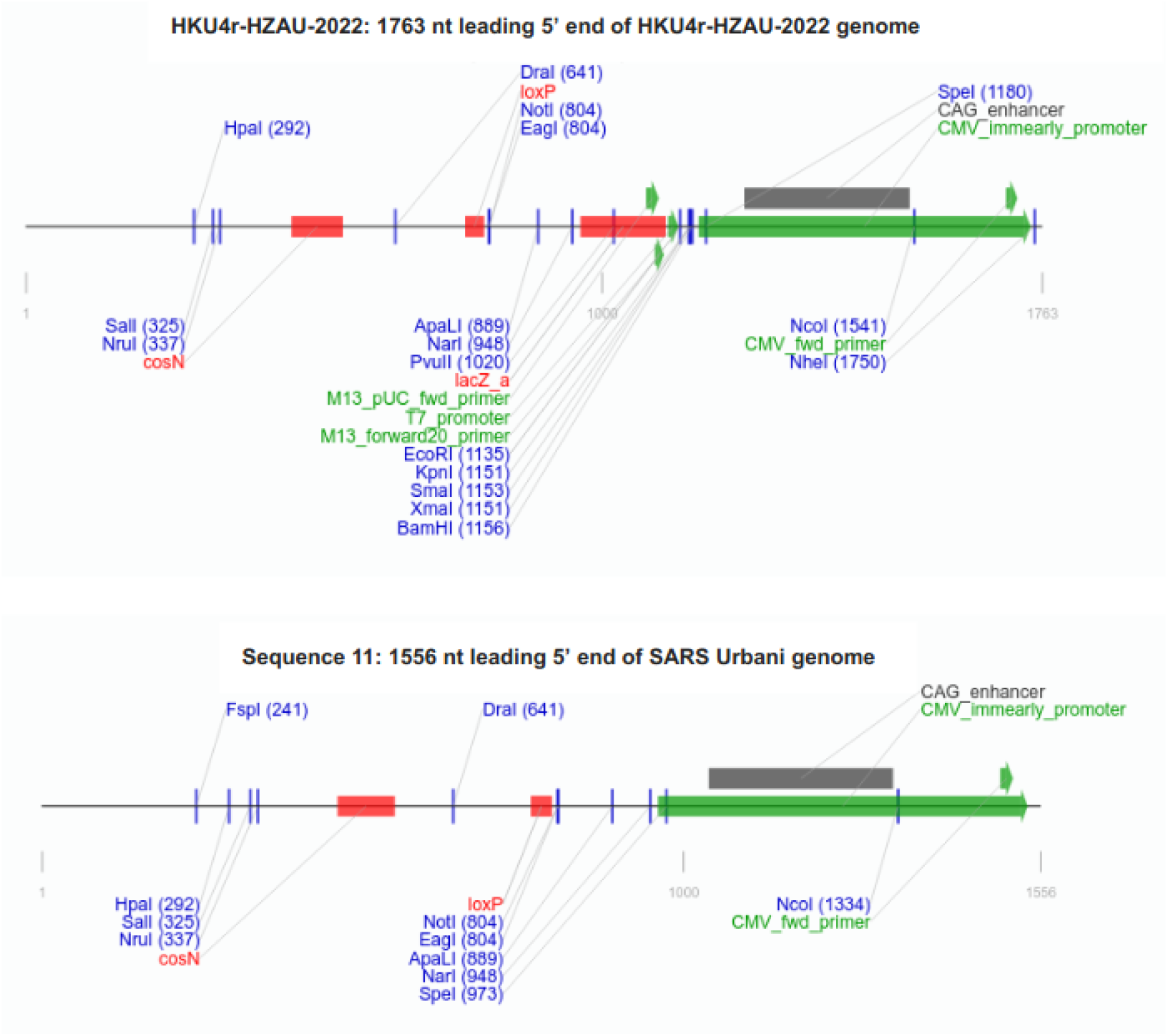
Comparison of the 1763 nt sequence of HKU4r-HZAU-2020 leading the HKU4r-HZAU-2020 genome and sequence upstream of the SARS genome from pBAC-SARS-CoV Sequence 11 from Patent WO2006236448 (accession CS480537.1).

**Supp. Fig. 5.**
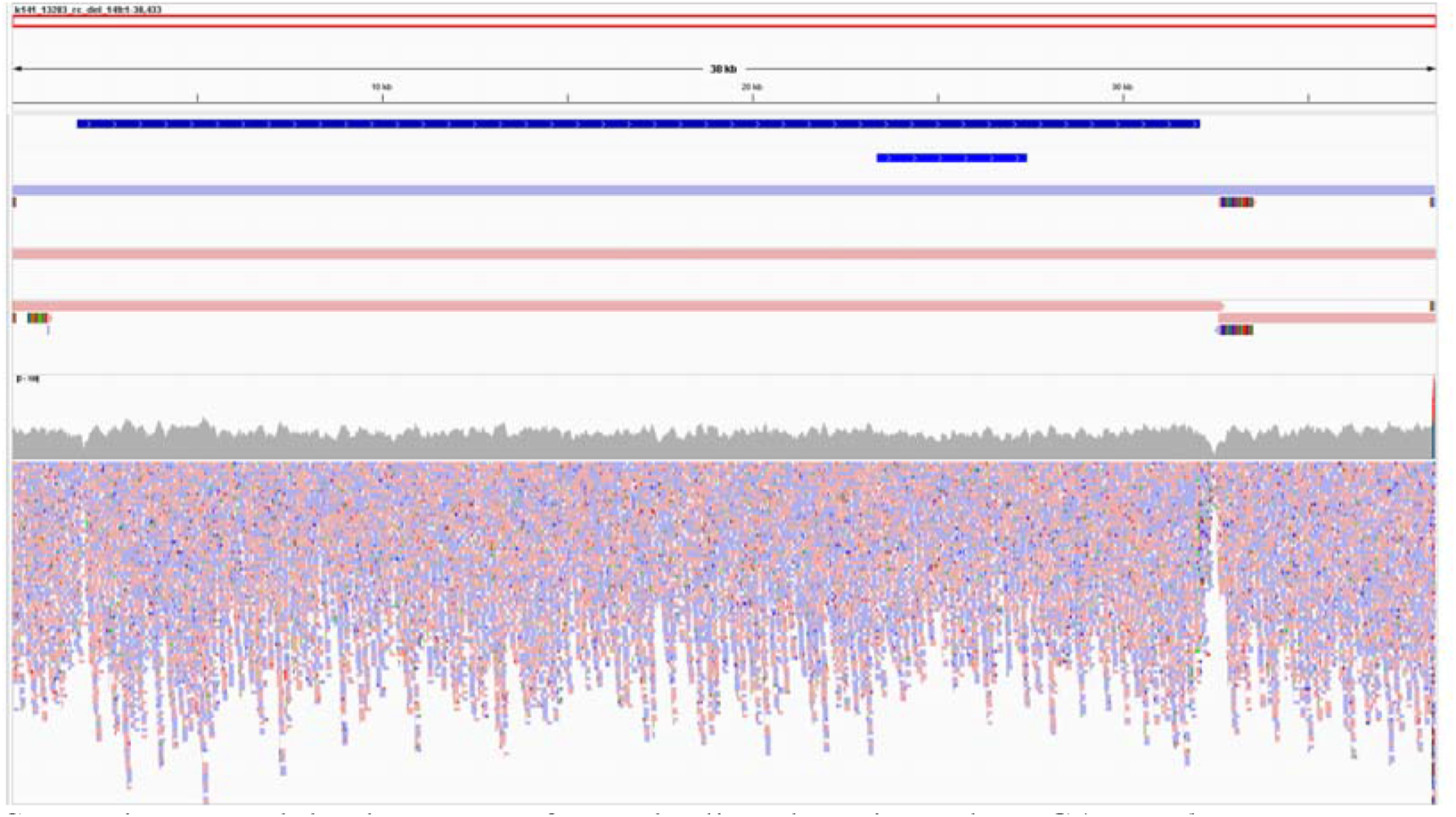
Read depth coverage for reads aligned to trimmed MEGAHIT *de novo* assembled contig k141_13283_del_149 (HKU4r-HZAU-2020 clone) for (top to bottom): HKU4r-HZAU-2020 genome (dark blue); b) S gene (blue); c) MEGAHIT assembled contig k141_13283_del_149; d) SPAdes assembled contig NODE_1_length_38592_cov_26.713566; e) CoronaSPAdes assembled contigs NODE_1_length_32683_cov_25.774854 and NODE_4_length_5879_cov_27.213917. Displayed in IGV.

**Supp. Fig. 6.**
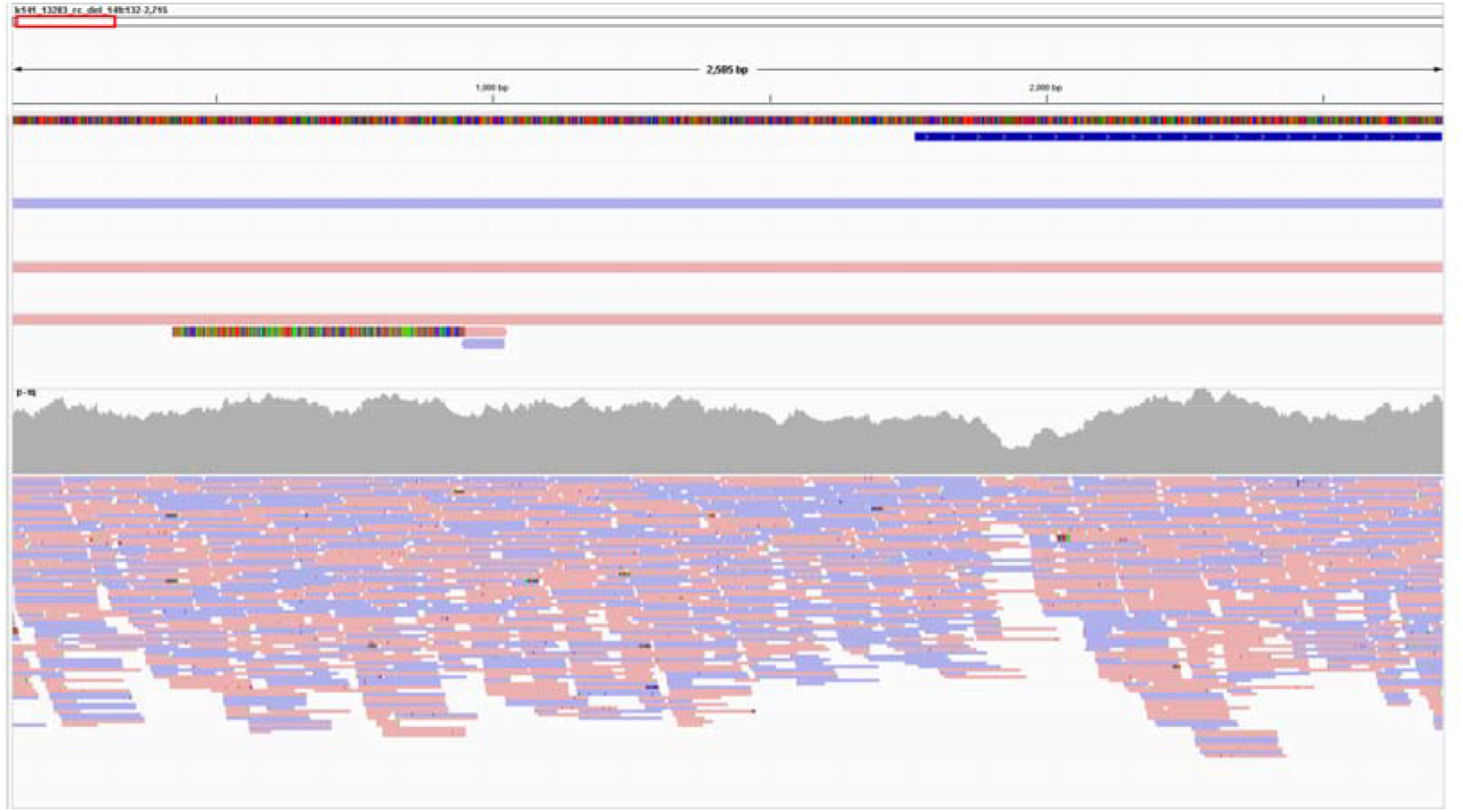
Read depth coverage for reads aligned to trimmed MEGAHIT *de novo* assembled contig k141_13283_del_149 (HKU4r-HZAU-2020 clone) zoomed in to 5’ end of the HKU4r-HZAU-2020 genome, for (top to bottom): HKU4r-HZAU-2020 genome (dark blue); b) MEGAHIT assembled contig k141_13283; c) SPAdes assembled contig NODE_1_length_38592_cov_26.713566; d) CoronaSPAdes assembled contigs NODE_1_length_32683_cov_25.774854 (two small CoronaSPAdes assembled contigs also shown). Displayed in IGV.

**Supp. Fig. 7.**
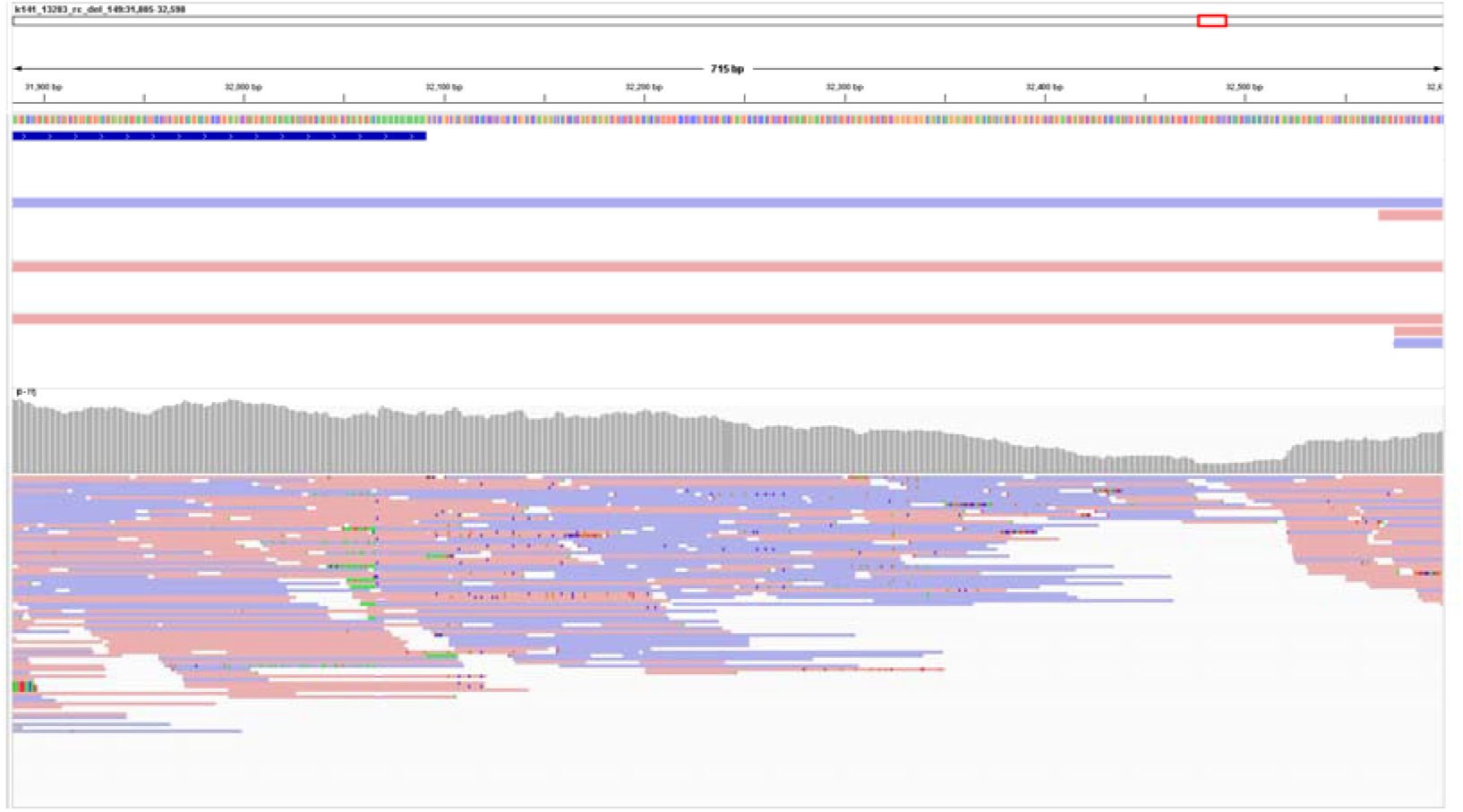
Read depth coverage for reads aligned to trimmed MEGAHIT *de novo* assembled contig k141_13283_del_149 (HKU4r-HZAU-2020 clone) zoomed in to 3’ end of the HKU4r-HZAU-2020 genome, for (top to bottom): HKU4r-HZAU-2020 genome (dark blue); b) MEGAHIT assembled contig k141_13283 (small section of contig k141_2146 also shown); c) SPAdes assembled contig NODE_1_length_38592_cov_26.713566; d) CoronaSPAdes assembled contig NODE_1_length_32683_cov_25.774854 (sections of contigs NODE_4_length_5879_cov_27.213917 and NODE_8159_length_935_cov_37.324826 also shown). Displayed in IGV.

**Supp. Fig. 8.**
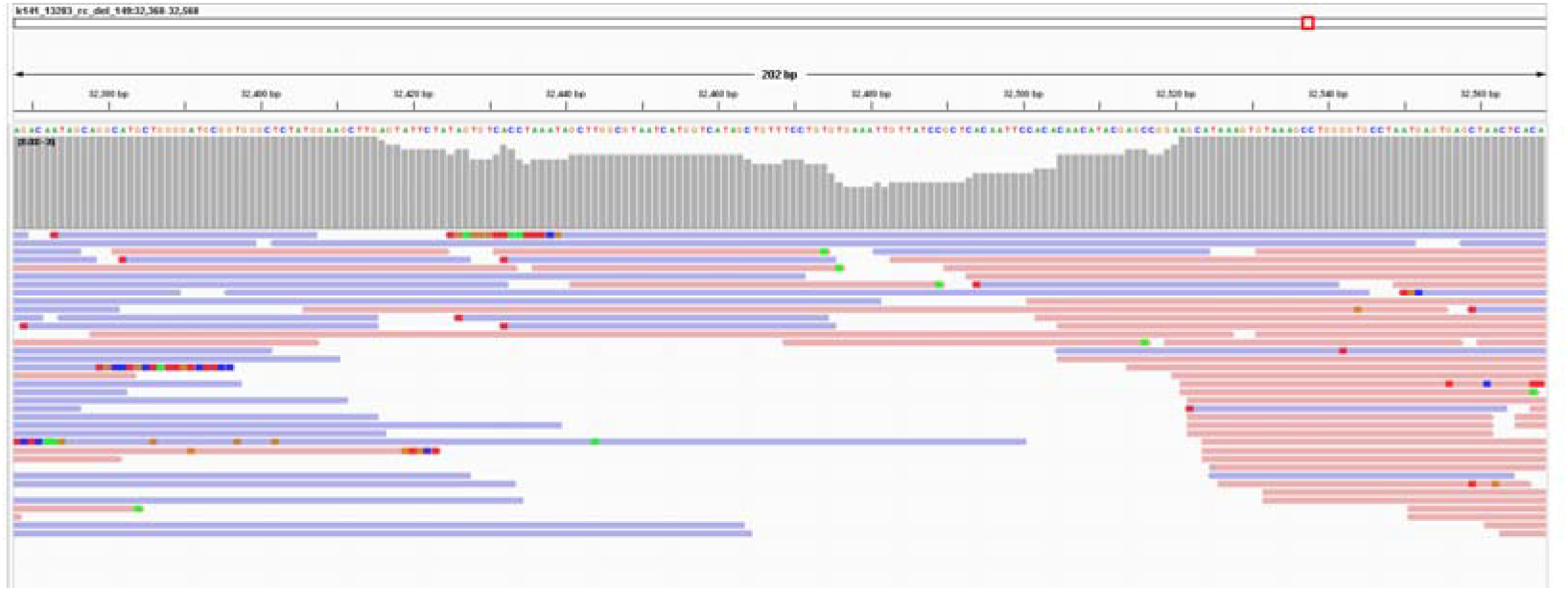
Low coverage region (9 to 19 reads) between 32,416 and 32,520 nt for four pooled SRA datasets aligned to the HKU4r-HZAU-2020 clone using minimap2. The low read depth region occurs within a vector sequence, immediately downstream of a bGH poly(A) signal. Read depth scale (gray) 0-20.

**Supp. Fig. 9.**
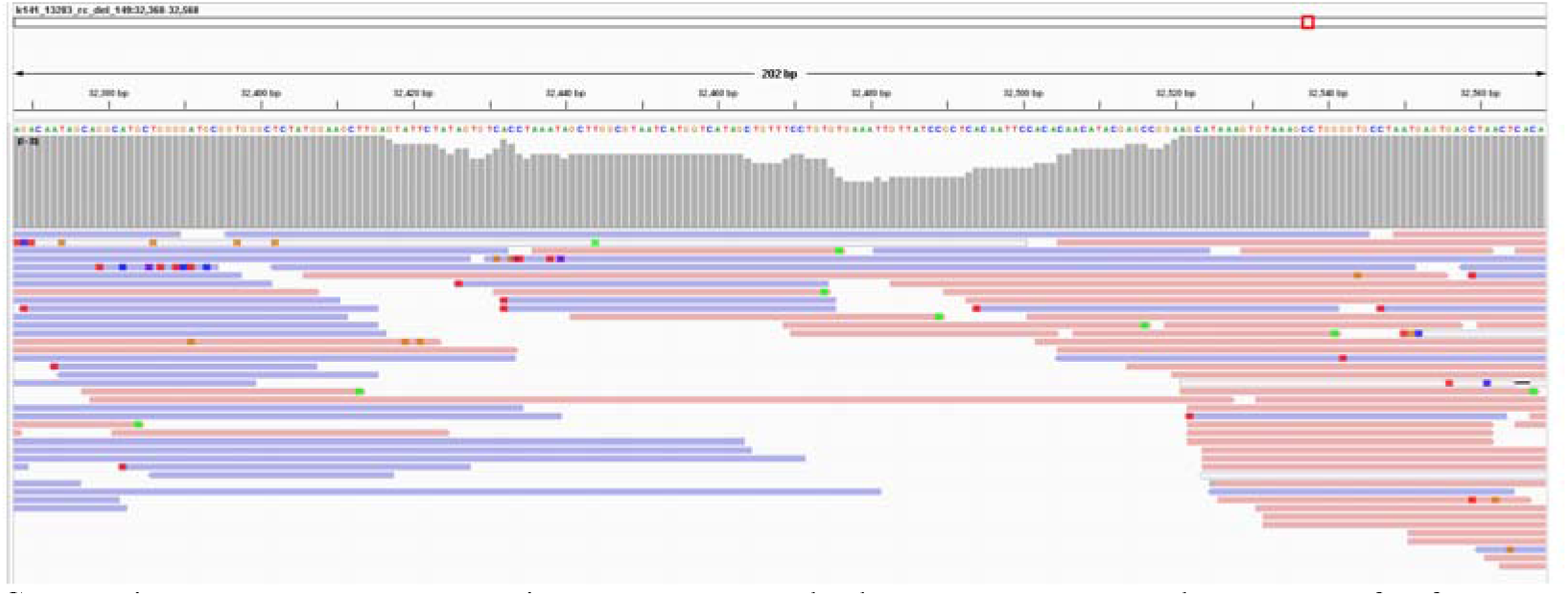
Low coverage region (10 to 19 reads) between 32,417 and 32,520 nt for four pooled SRA datasets aligned to the HKU4r-HZAU-2020 clone using bowtie2. Read depth scale (gray) 0-20.

**Supp. Fig. 10.**
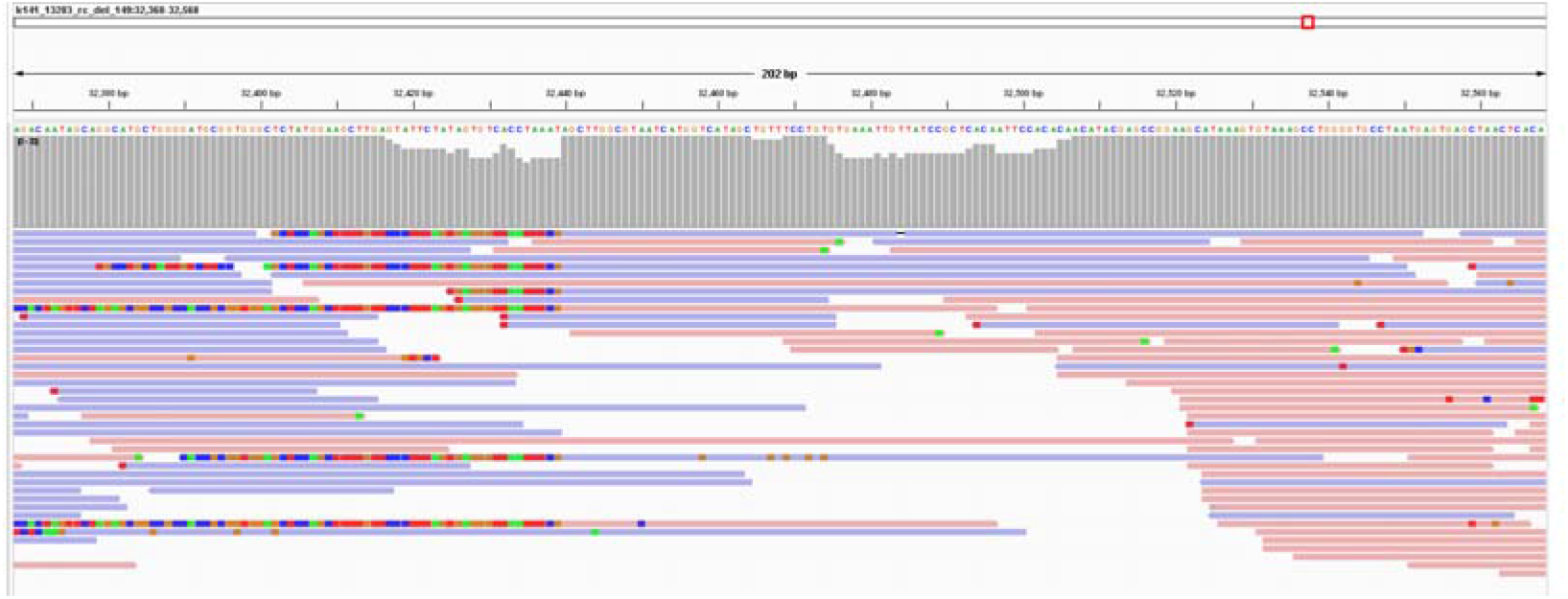
Low coverage region (14 to 19 reads) between 32,417 and 32,520 nt for four pooled SRA datasets aligned to the HKU4r-HZAU-2020 clone using bwa-mem2. Read depth scale (gray) 0-20.

**Supp. Fig. 11.**
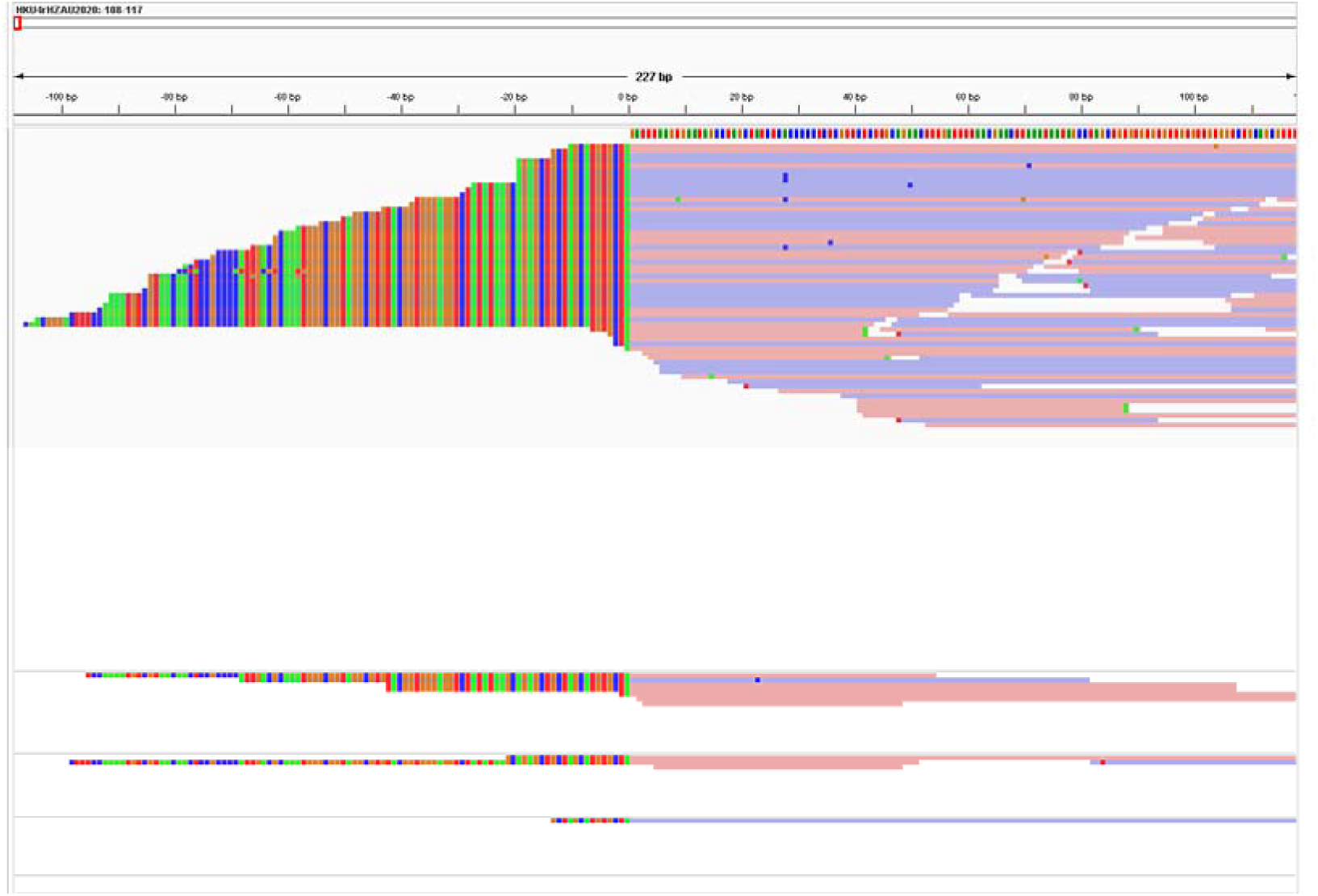
Reads mapping to the HKU4r-HZAU-2020 genome show soft-clipped ends upstream of the 5’ end of the CoV genome. Top to bottom: SRR10915173; SRR10915174; SRR10915168 and SRR10915167. Mapped using minimap2, displayed using IGV.

**Supp. Fig. 12.**
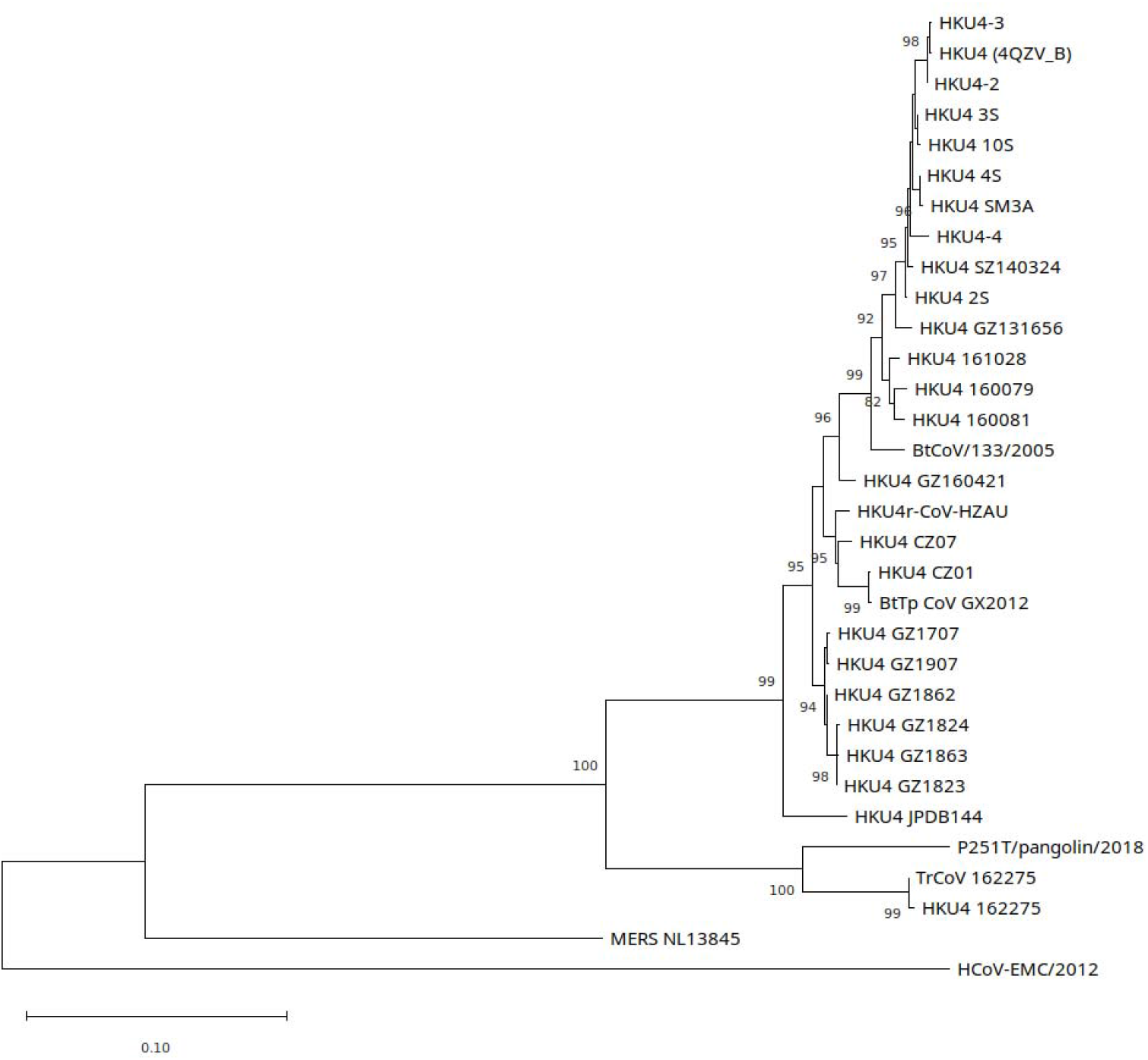
Maximum likelihood phylogenetic trees for S protein for selected HKU4-related CoVs and MERS CoVs, generated using a WAG+R+F model in PhyML using smart model selection (Lefort et al., 2017). Tree rooted on midpoint. Branch support values less than 80 are hidden. See Supp. Info. 1.7 for GenBank accession numbers.

**Supp. Fig. 13.**
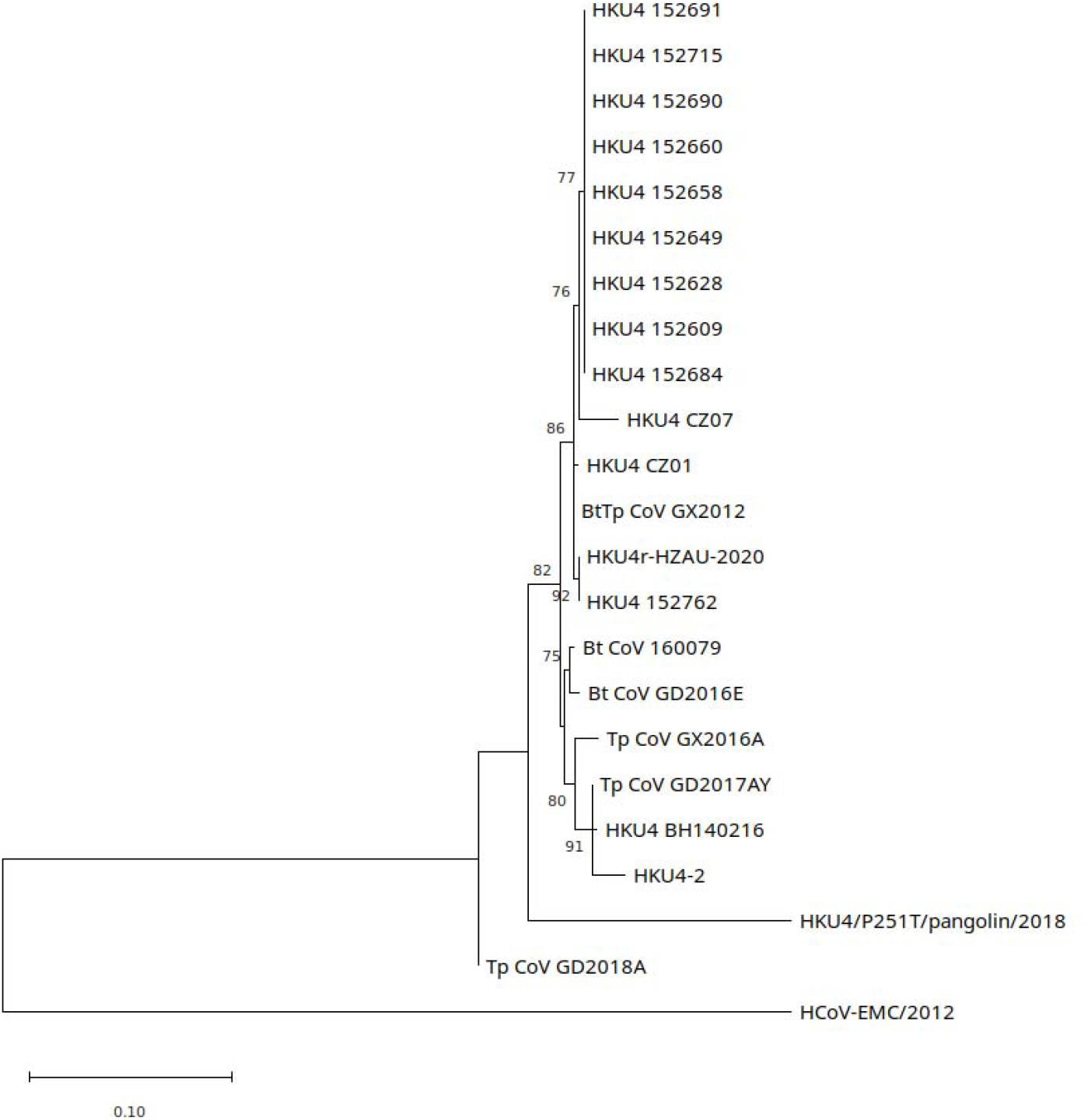
Maximum likelihood phylogenetic tree for a 399 nt partial RdRp segment. Generated using a TN93+G model in PhyML using smart model selection (Lefort et al., 2017). Tree rooted on midpoint. See Supp. Info. 1.4 for Genbank accession numbers.

**Supp. Fig. 14.**
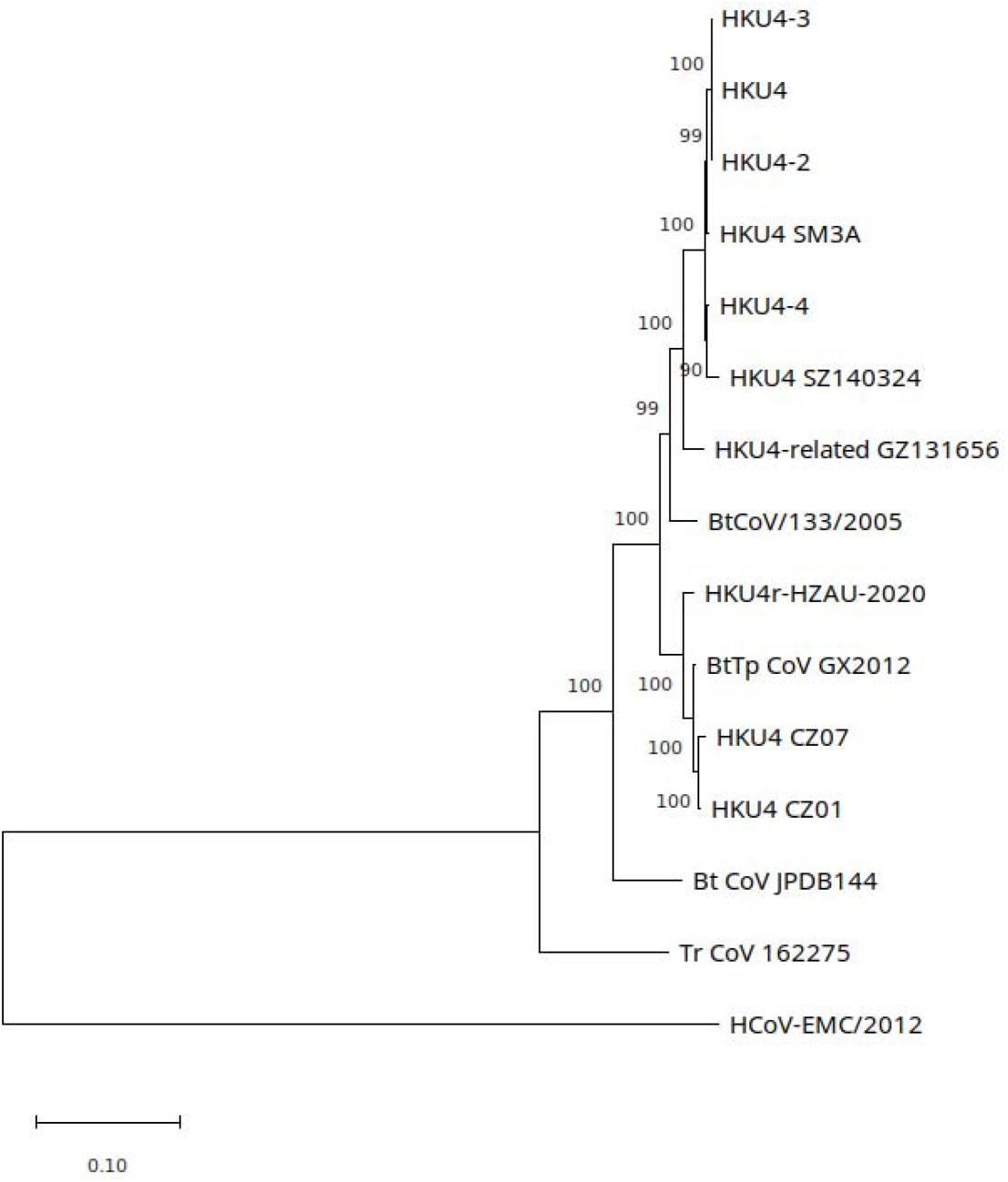
Full genome maximum likelihood phylogenetic tree generated using a GTR+G+I model in MEGA11 with 1000 bootstrap replicates. Tree rooted on midpoint. See Supp. Info. 1.4 for Genbank accession numbers.

**Supp. Fig. 15.**
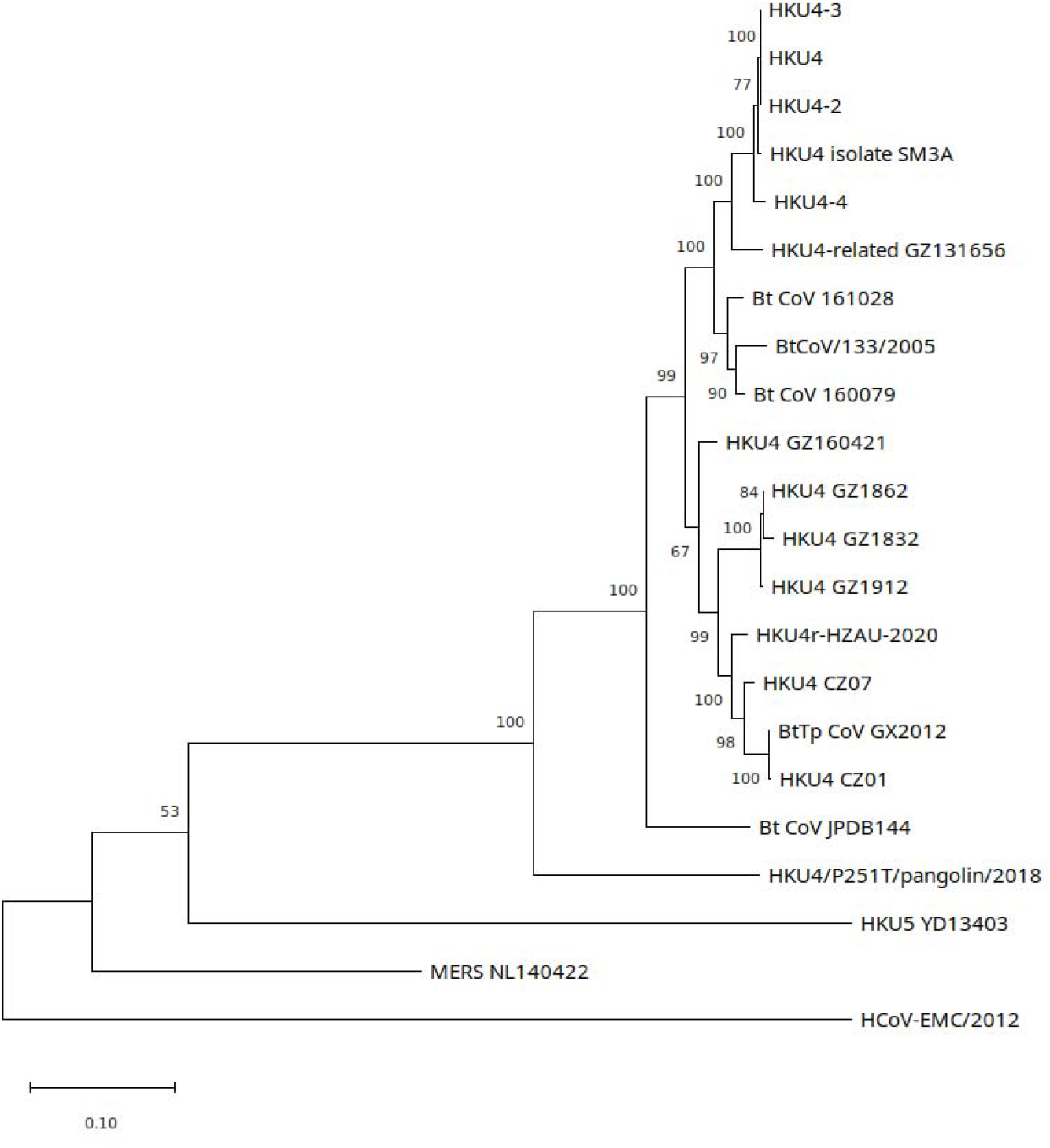
Maximum likelihood phylogenetic trees for S gene for selected HKU4-related CoVs, MERS CoVs, and HKU5 isolate YD13403, generated using a GTR+G model in MEGA11 with 1000 bootstrap replicates. Tree rooted on midpoint. See Supp. Info. 1.4 for Genbank accession numbers.

**Supp. Fig. 16.**
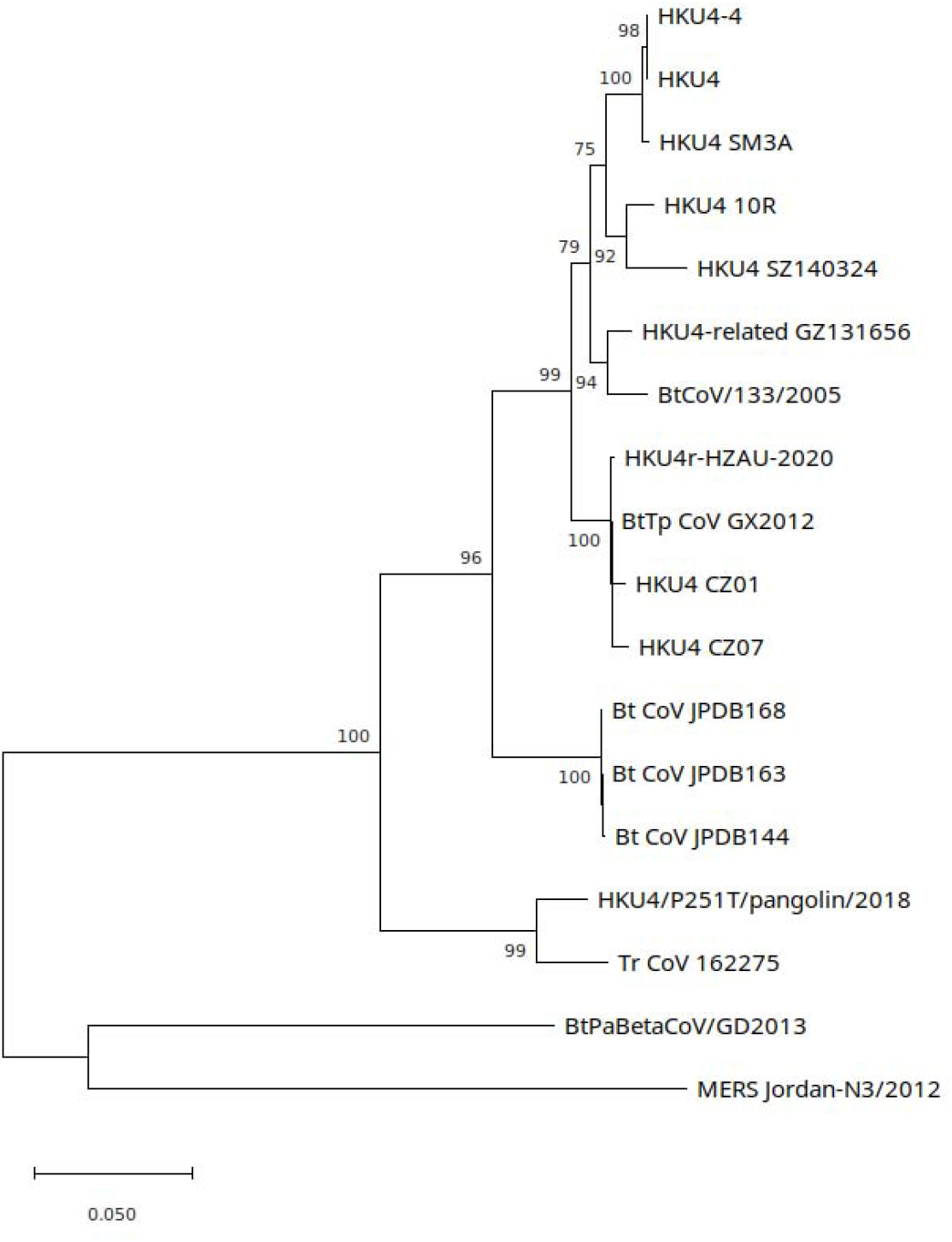
RdRp maximum likelihood phylogenetic tree, generated using a TN93+G model in MEGA11 with 100 bootstrap replicates. Tree rooted on midpoint. See Supp. Info. 1.4 for Genbank accession numbers.

**Supp. Fig. 17.**
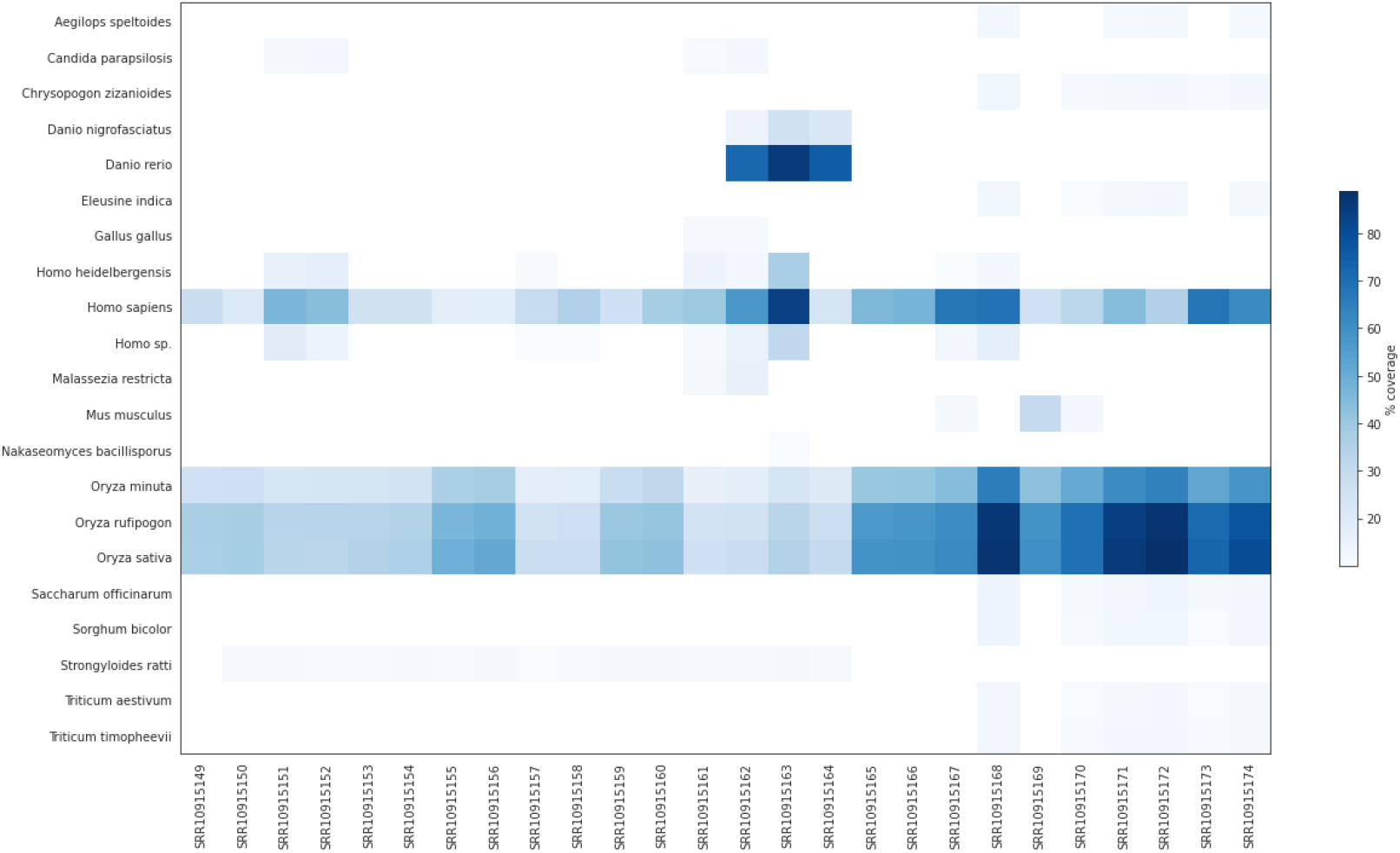
Mitochondrial genome coverage using a minimum coverage cutoff of 10% in all SRA datasets in BioProject PRJNA602160. See Supp. Info. 1.1 for SRA dataset descriptions.

**Supp. Fig. 18.**
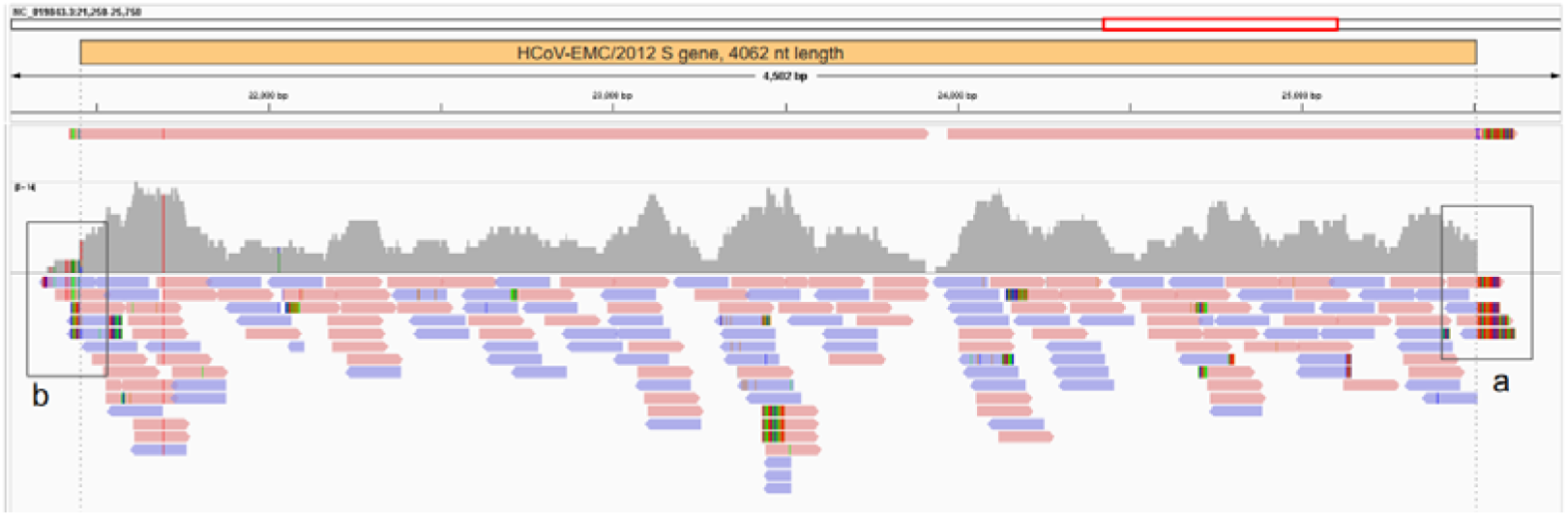
MERS-CoV genome alignment. In top track pooled datasets SRR10915167, SRR10915168, SRR10915173 and SRR10915174 were *de novo* assembled using MEGAHIT and contigs aligned to MERS-CoV reference genome HCoV-EMC/2012 (NC_019843.3). Middle track shows pooled read depth, scale 0-14. Bottom track shows pooled read alignments using minimap2. Boxes drawn around 5’ end (b) and 3’ end (a) of S gene and shown in Supp. Figs. 19-20.

**Supp. Fig. 19.**
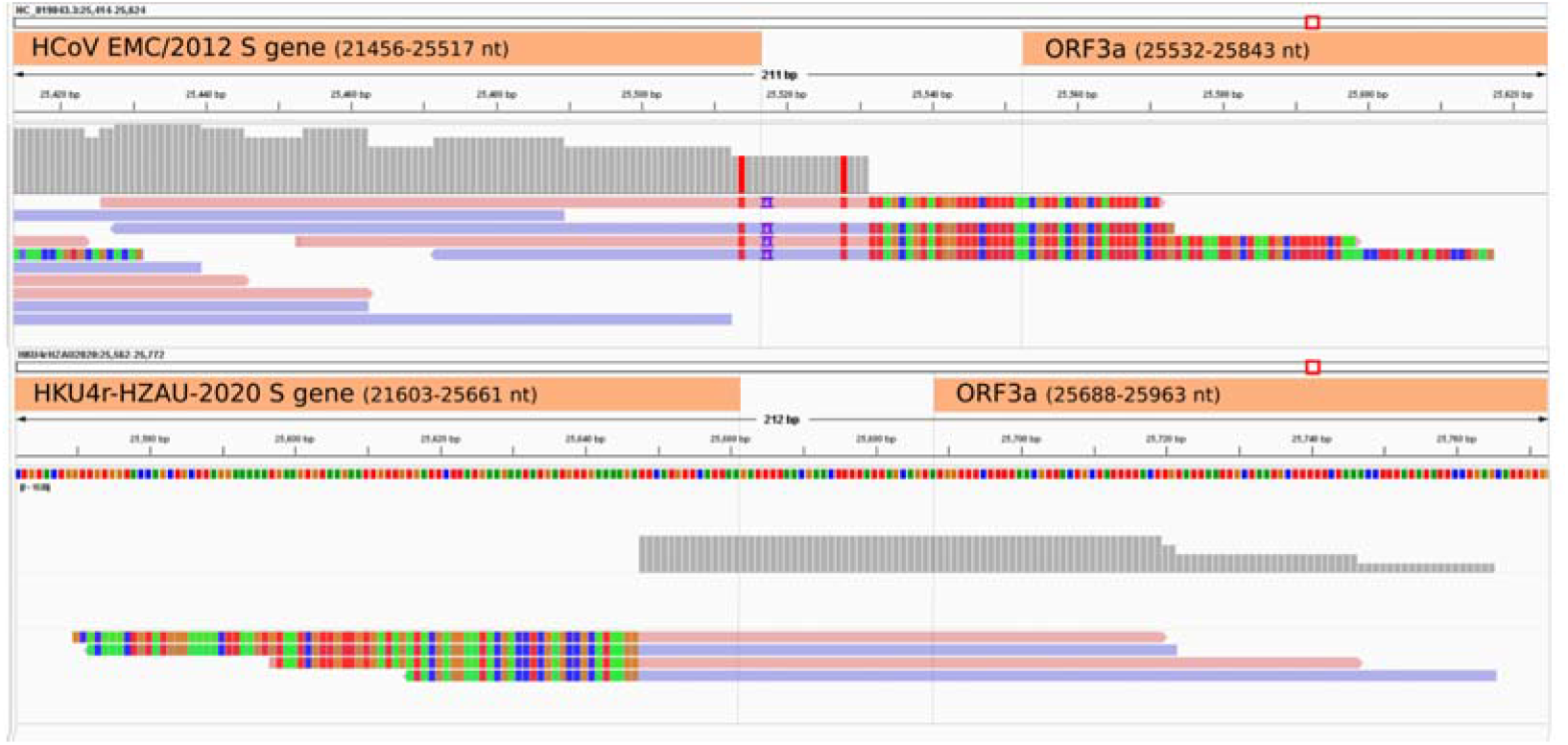
MERS-CoV and HKU4r-CoV-2020 3’ end of S gene alignments. Upper figure: four reads aligning to the 3’ end of the HCoV-EMC/2012 S gene exhibit a mismatch with the non-coding region and ORF3a gene downstream of the S gene. Lower figure: The same four reads align perfectly with the last 14 nt of the HKU4r-HZAU-2020 S gene sequence and downstream region. Reads mapped using bowtie2 using --local parameter setting. Displayed using IGV. See Supp. Data for read statistics.

**Supp. Fig. 20.**
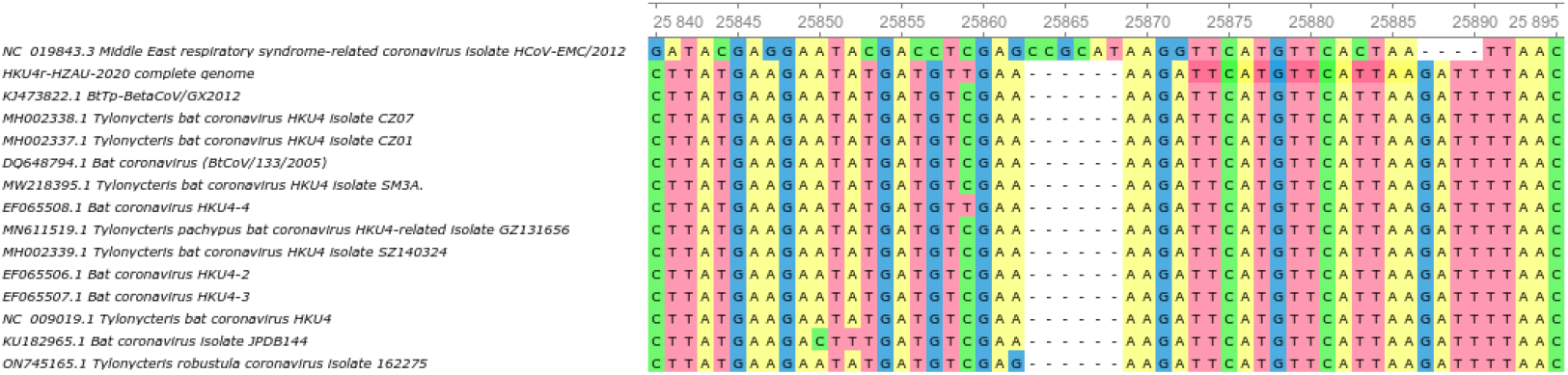
Multiple sequence alignment showing 3’ end of the S gene region for MERS-CoV HCoV-EMC/2012 and selected HKU-related CoVs. The last 14 nt of the S gene is positioned at 25873-25886 nt (multiple sequence reference numbers). This section is conserved for all sequences except for T25883C (T25514C in MERS-CoV reference) in HCoV-EMC/2012.

**Supp. Fig. 21.**
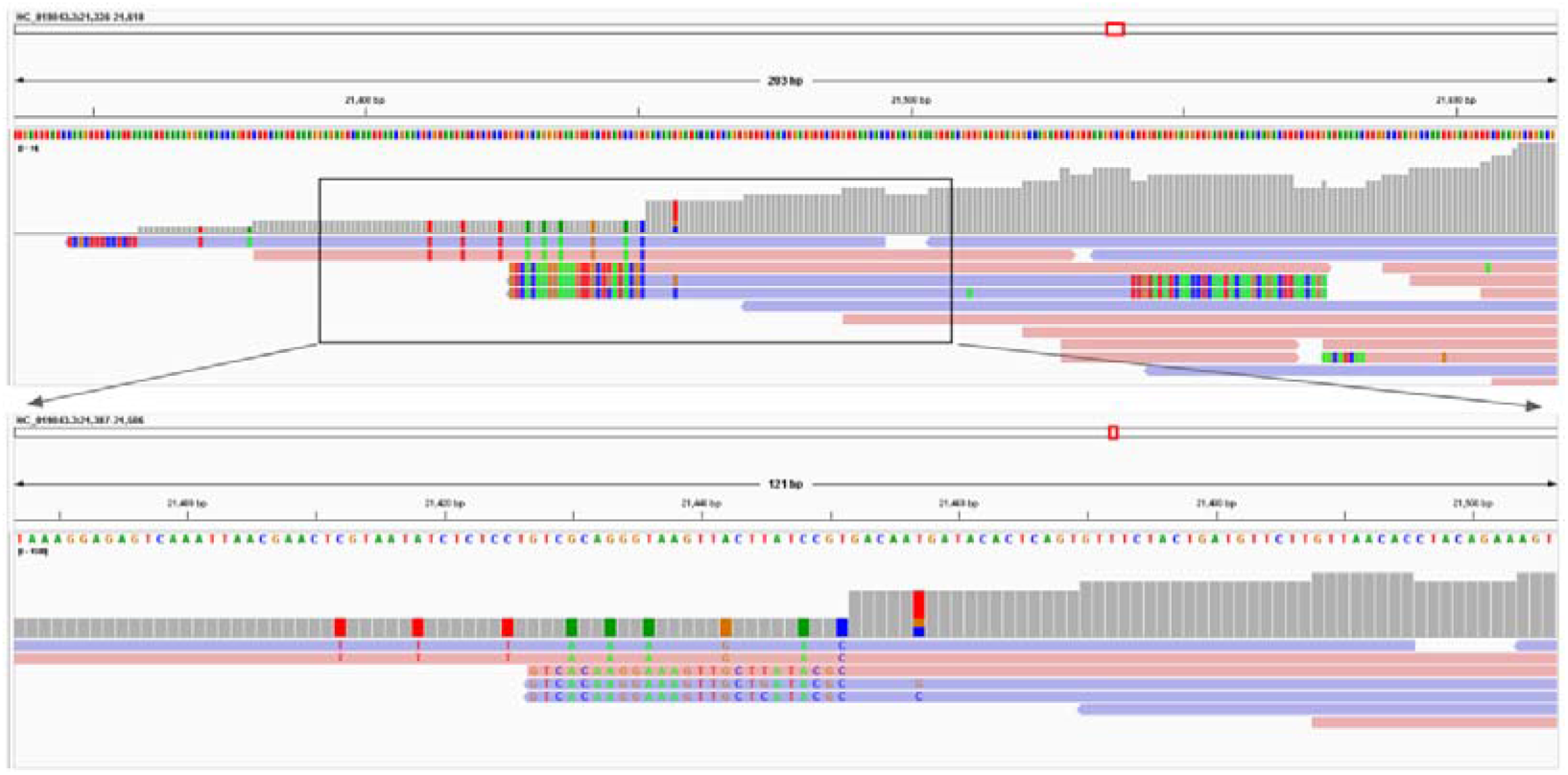
MERS-CoV genome alignment. Pooled datasets SRR10915167, SRR10915168, SRR10915173 and SRR10915174 were aligned to the MERS-CoV reference genome HCoV-EMC/2012 (NC_019843.3) using minimap2. The 5’ end of the S gene is located at position 21a,456 which is located 4 nt downstream of a region with numerous SNV’s relative to HCoV-EMC/2012. Note three reads with apparent 5’ soft clip ends have (only) either 6 or 7 SNV’s for a 19/25 or 18/25 nt match in this region to HKU4r-HZAU-2020. See Supp. Data for read statistics.

**Supp. Fig. 21.**
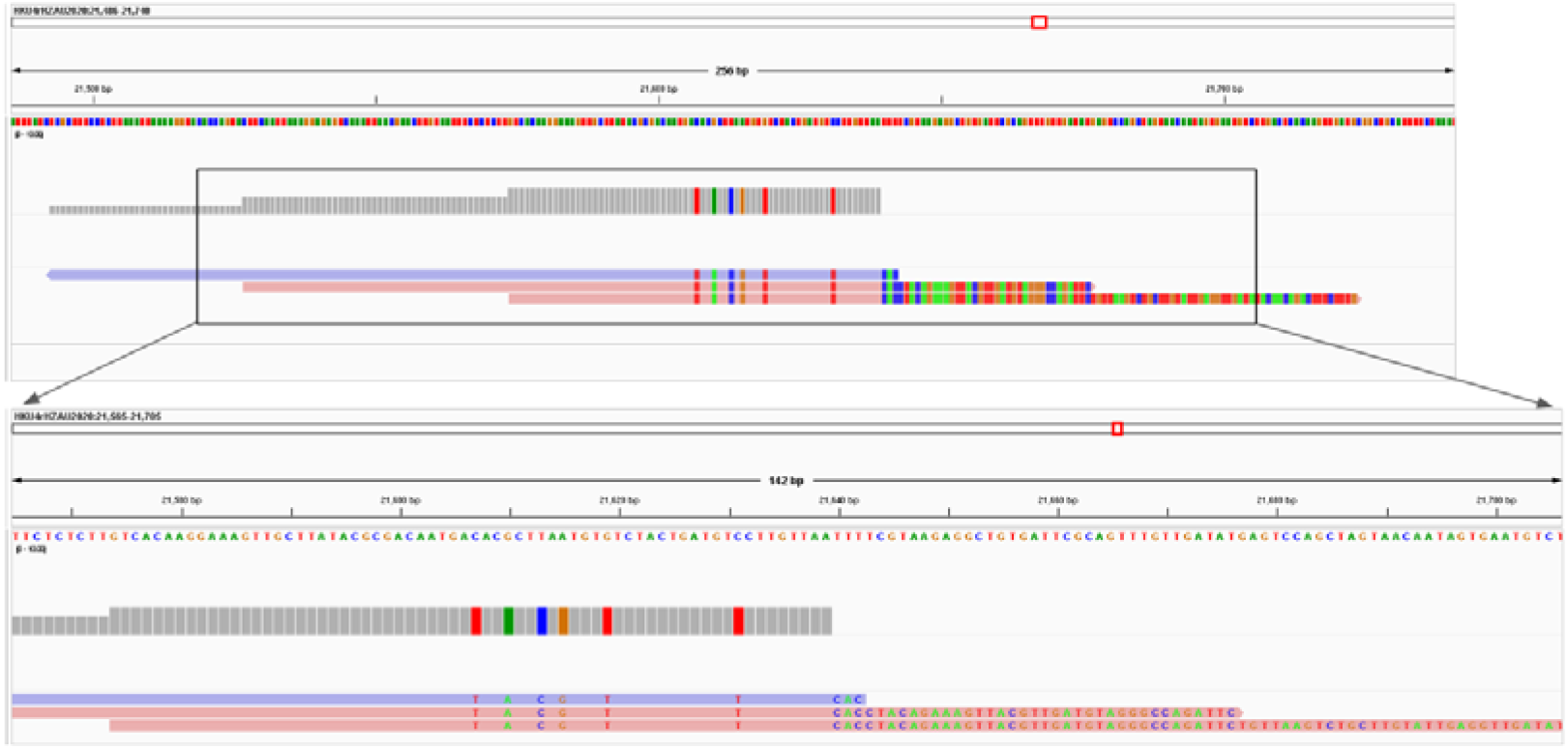
Three reads covering the 5’ end of the MERS-CoV S gene shown in Supp. Fig. 20 were aligned to the HKU4r-HAZU-2020 genome sequence using minimap2. The reads exhibit a 100% identity to HKU4r-HZAU-2020 at their 5’ ends.

**Supp. Fig. 21.**
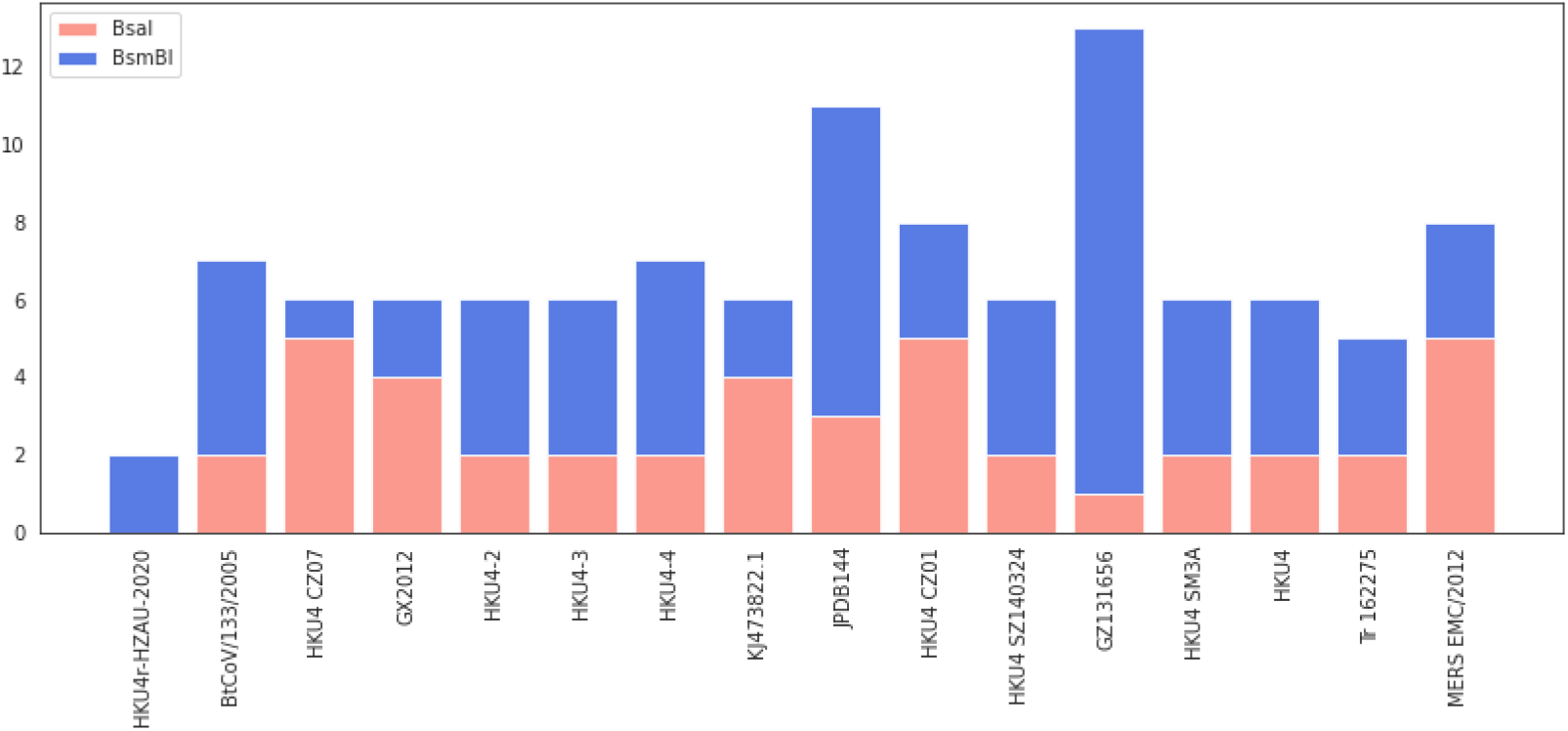
BsaI and BsmBI restriction enzyme recognition site counts in the HKU4r-HZAU-2020 genome, compared with the 14 closest related HKU4-related CoVs, MERS HCoV EMC/2012.

**Supp. Fig. 22.**
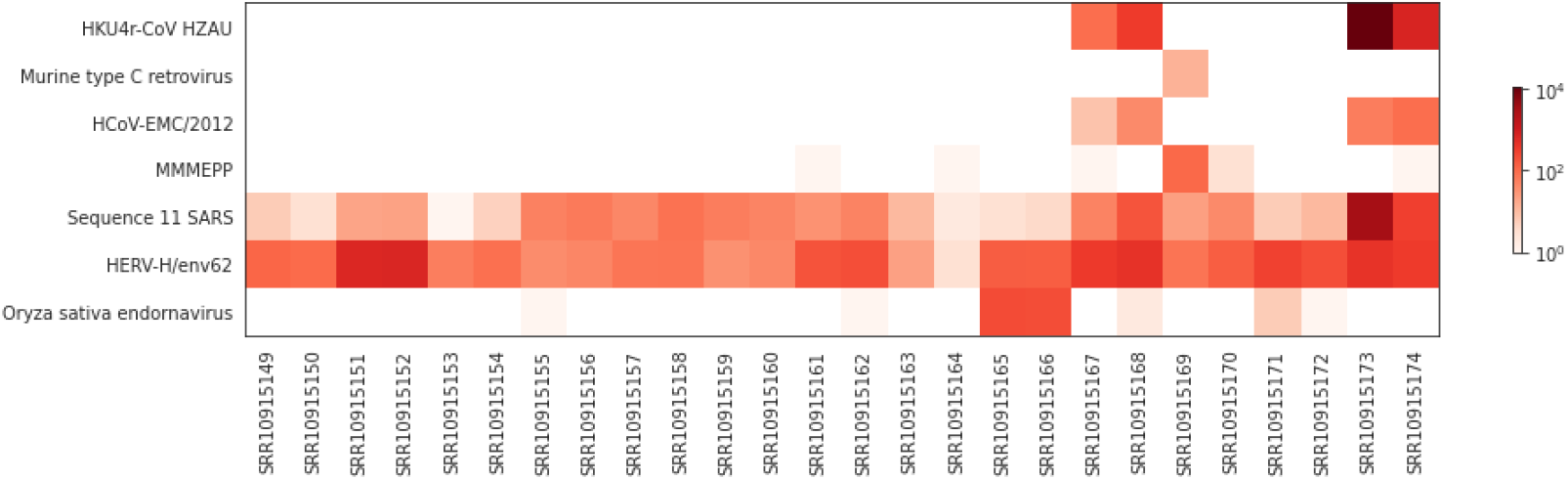
Number of reads mapping to virus genome sequences and Sequence 11 synthetic construct with greater than 10% coverage identified in BioProject PRJNA602160.

**Supp. Table 1.**
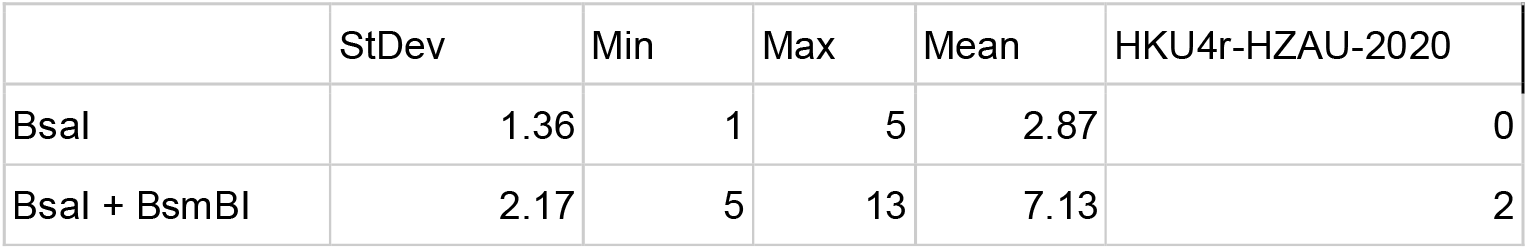
BsaI and BsmBI restriction site basic statistics for 14 HKU4-related CoV genomes and HKU4r-HZAU-2020.

## References

Addgene. Addgene: Analyze Sequence. 2022. https://www.addgene.org/analyze-sequence/

Almazán F, DeDiego ML, Galán C, et al. Construction of a Severe Acute Respiratory Syndrome Coronavirus Infectious cDNA Clone and a Replicon To Study Coronavirus RNA Synthesis. J Virol. 2006;80(21):10900–10906. doi:10.1128/JVI.00385-06

Almazán F, Galán C, Enjuanes L. Engineering Infectious cDNAs of Coronavirus as Bacterial Artificial Chromosomes. In: Coronaviruses: Methods and Protocols. Vol 454.; 2008:275–291. doi:10.1007/978-1-59745-181-9_20

Alraddadi BM, Watson JT, Almarashi A, et al. Risk Factors for Primary Middle East Respiratory Syndrome Coronavirus Illness in Humans, Saudi Arabia, 2014. Emerg Infect Dis. 2016;22(1):49–55. doi:10.3201/eid2201.151340

Andersen KG, Rambaut A, Lipkin WI, Holmes EC, Garry RF. The proximal origin of SARS-CoV-2. Nat Med. 2020;26(4):450–452. doi:10.1038/s41591-020-0820-9

Agarwala R, Barrett T, Beck J, et al. Database resources of the National Center for Biotechnology Information. Nucleic Acids Res. 2018;46(D1):D8–D13. doi:10.1093/nar/gkx1095

Ballenghien M, Faivre N, Galtier N. Patterns of cross-contamination in a multispecies population genomic project: detection, quantification, impact, and solutions. BMC Biol. 2017;15(1):25. doi:10.1186/s12915-017-0366-6

Bamdst. https://github.com/shiquan/bamdst. Downloaded 2021-7-05.

Belouzard S, Chu VC, Whittaker GR. Activation of the SARS coronavirus spike protein via sequential proteolytic cleavage at two distinct sites. Proc Natl Acad Sci U S A. 2009;106(14):5871–5876. doi:10.1073/pnas.0809524106

Bermingham A, Chand MA, Brown CS, et al. Severe respiratory illness caused by a novel coronavirus, in a patient transferred to the United Kingdom from the Middle East, September 2012. Eurosurveillance. 2012;17(40). doi:10.2807/ese.17.40.20290-en

Boni MF, Posada D, Feldman MW. An Exact Nonparametric Method for Inferring Mosaic Structure in Sequence Triplets. Genetics. 2007;176(2):1035–1047. doi:10.1534/genetics.106.068874

Butler D. Engineered bat virus stirs debate over risky research. Nature. 2015;(November):1–2. doi:10.1038/nature.2015.18787

Cantalupo PG, Pipas JM. Detecting viral sequences in NGS data. Curr Opin Virol. 2019;39:41–48. doi:10.1016/j.coviro.2019.07.010

Chen L, Yu B, Hua J, et al. Construction of a full-length infectious bacterial artificial chromosome clone of duck enteritis virus vaccine strain. Virol J. 2013;10(1):1. doi:10.1186/1743-422X-10-328

Chen S, Zhou Y, Chen Y, Gu J. fastp: an ultra-fast all-in-one FASTQ preprocessor. Bioinformatics. 2018;34(17):i884–i890. doi:10.1093/bioinformatics/bty560

Chen S, He C, Li Y, Li Z, Melançon CE. A computational toolset for rapid identification of SARS-CoV-2, other viruses and microorganisms from sequencing data. Brief Bioinform. 2020;00(August):1–12. doi:10.1093/bib/bbaa231

Chen Z, Bao L, Chen C, et al. Human Neutralizing Monoclonal Antibody Inhibition of Middle East Respiratory Syndrome Coronavirus Replication in the Common Marmoset. J Infect Dis. 2017;215(12):1807–1815. doi:10.1093/infdis/jix209

Cohen J. Wuhan coronavirus hunter Shi Zhengli speaks out. 2020. In Science (Vol. 369, Issue 6503, pp. 487–488). American Association for the Advancement of Science. https://doi.org/10.1126/science.369.6503.487

Csabai I, Stéger J. Unique SARS-CoV-2 variant found in public sequence data of Antarctic soil samples collected in 2018-2019. Published online 2021. doi:10.13140/RG.2.2.30183.37281

Daszak P. Understanding the Risk of Bat Coronavirus Emergence. 2021. NIH Grant: 5R01AI110964-05. https://www.nih.gov/sites/default/files/institutes/foia/20211020-risk-of-bat-emergence.pdf

Davis MW, Jorgensen EM. ApE, A Plasmid Editor: A Freely Available DNA Manipulation and Visualization Program. Front Bioinforma. 2022;2(February):1–15. doi:10.3389/fbinf.2022.818619

El-Kafrawy SA, Corman VM, Tolah AM, et al. Enzootic patterns of Middle East respiratory syndrome coronavirus in imported African and local Arabian dromedary camels: a prospective genomic study. Lancet Planet Heal. 2019;3(12):e521–e528. doi:10.1016/S2542-5196(19)30243-8

Engler C, Kandzia R, Marillonnet S. A one pot, one step, precision cloning method with high throughput capability. PLoS One. 2008;3(11). doi:10.1371/journal.pone.0003647

Engler C, Marillonnet S. Golden Gate cloning. Methods Mol Biol. 2014;1116:119–131. doi:10.1007/978-1-62703-764-8_9

Enjuanes L, DeDiego ML, Álvarez E, Almazán F. Attenuated SARS and use as a vaccine. 2006. https://patentscope.wipo.int/search/en/detail.jsf?docId=WO2006136448

Enjuanes L, DeDiego ML, Álvarez E, Deming D, Sheahan T, Baric R. Vaccines to prevent severe acute respiratory syndrome coronavirus-induced disease. Virus Res. 2008;133(1):45–62. doi:10.1016/j.virusres.2007.01.021

Epstein JH, Anthony SJ, Islam A, et al. Nipah virus dynamics in bats and implications for spillover to humans. Proc Natl Acad Sci U S A. 2020;117(46):29190–29201. doi:10.1073/pnas.2000429117

Fan Y, Zhao K, Shi Z-L, Zhou P. Bat Coronaviruses in China. Viruses. 2019;11(3):210. doi:10.3390/v11030210

Fehr AR, Perlman S. Coronaviruses: An overview of their replication and pathogenesis. In: Coronaviruses: Methods and Protocols. Springer New York; 2015:1–23. doi:10.1007/978-1-4939-2438-7_1

Gibbs MJ, Armstrong JS, Gibbs AJ. Sister-Scanning: a Monte Carlo procedure for assessing signals in recombinant sequences. Bioinformatics. 2000;16(7):573–582. doi:10.1093/bioinformatics/16.7.573

Guindon S, Dufayard J-F, Lefort V, Anisimova M, Hordijk W, Gascuel O. New Algorithms and Methods to Estimate Maximum-Likelihood Phylogenies: Assessing the Performance of PhyML 3.0. Syst Biol. 2010;59(3):307–321. doi:10.1093/sysbio/syq010

Han X, Qi J, Song H, et al. Structure of the S1 subunit C-terminal domain from bat– derived coronavirus HKU5 spike protein. Virology. 2017;507(January):101–109. doi:10.1016/j.virol.2017.04.016

Heckert RA, Kozlovac JP. Special Considerations for Agriculture Pathogen Biosafety. In: Biological Safety. ASM Press; 2014:579–586. doi:10.1128/9781555815899.ch32

Holmes EC, Worobey M, Rambaut A. Phylogenetic evidence for recombination in dengue virus. Mol Biol Evol. 1999;16(3):405–409. doi:10.1093/oxfordjournals.molbev.a026121

Hu B, Zeng L-P, Yang X-L, et al. Discovery of a rich gene pool of bat SARS-related coronaviruses provides new insights into the origin of SARS coronavirus. Drosten C, ed. PLOS Pathog. 2017;13(11):e1006698. doi:10.1371/journal.ppat.1006698

Huang Y, Huang C-X, Wang W-F, Liu H, Wang H-L. Zebrafish miR-462-731 is required for digestive organ development. Comp Biochem Physiol Part D Genomics Proteomics. 2020;34:100679. doi:10.1016/j.cbd.2020.100679

Inayoshi Y, Oguro S, Tanahashi E, et al. Bacterial artificial chromosome-based reverse genetics system for cloning and manipulation of the full-length genome of infectious bronchitis virus. Curr Res Microb Sci. 2022;3(July):100155. doi:10.1016/j.crmicr.2022.100155

Islam R, Raju RS, Tasnim N, et al. Choice of assemblers has a critical impact on de novo assembly of SARS-CoV-2 genome and characterizing variants. Brief Bioinform. 2021;22(5):1–11. doi:10.1093/bib/bbab102

Johnson M, Zaretskaya I, Raytselis Y, Merezhuk Y, McGinnis S, Madden TL. NCBI BLAST: a better web interface. Nucleic Acids Res. 2008;36(Web Server):W5–W9. doi:10.1093/nar/gkn201

Jones A, Massey SE, Zhang D, Deigin Y, Quay SC. Forensic Analysis of Novel SARS2r-CoV Identified in Game Animal Datasets in China Shows Evolutionary Relationship to Pangolin GX CoV Clade and Apparent Genetic Experimentation. Appl Microbiol. 2022;2(4):882–904. doi:10.3390/applmicrobiol2040068

Katoh K, Standley DM. MAFFT multiple sequence alignment software version 7: Improvements in performance and usability. Mol Biol Evol. 2013;30(4):772–780. doi:10.1093/molbev/mst010

Katz, K. S., Shutov, O., Lapoint, R., Kimelman, M., Brister, J. R., & O’Sullivan, C. (2021). STAT: a fast, scalable, MinHash-based k-mer tool to assess Sequence Read Archive next-generation sequence submissions. Genome Biology, 22(1), 270. https://doi.org/10.1186/s13059-021-02490-0

Klein MR. Classification of biological agents. National Institute for Public Health and the Environment. RIVM Letter report 205084002/2012 https://www.rivm.nl/bibliotheek/rapporten/205084002.pdf

Kleine-Weber H, Elzayat MT, Hoffmann M, Pöhlmann S. Functional analysis of potential cleavage sites in the MERS-coronavirus spike protein. Sci Rep. 2018;8(1):16597. doi:10.1038/s41598-018-34859-w

Klotz L. Human error in high-biocontainment labs: a likely pandemic threat. Bulletin of the Atomic Scientists. February 25, 2019. https://thebulletin.org/2019/02/human-error-in-high-biocontainment-labs-a-likely-pandemic-threat/

Langmead B, Salzberg SL. Fast gapped-read alignment with Bowtie 2. Nat Methods. 2012;9(4):357–359. doi:10.1038/nmeth.1923

Latinne A, Hu B, Olival KJ, et al. Origin and cross-species transmission of bat coronaviruses in China. Nat Commun. 2020;11(1):4235. doi:10.1038/s41467-020-17687-3

Lau SKP, Fan RYY, Luk HKH, et al. Replication of MERS and SARS coronaviruses in bat cells offers insights to their ancestral origins. Emerg Microbes Infect. 2018;7(1):1–11. doi:10.1038/s41426-018-0208-9

Lau SKP, Fan RYY, Zhu L, et al. Isolation of MERS-related coronavirus from lesser bamboo bats that uses DPP4 and infects human-DPP4-transgenic mice. Nat Commun. 2021;12(1):216. doi:10.1038/s41467-020-20458-9

Lefort V, Longueville J-E, Gascuel O. SMS: Smart Model Selection in PhyML. Mol Biol Evol. 2017;34(9):2422–2424. doi:10.1093/molbev/msx149

Lewis G, Jordan JL, Relman DA, et al. The biosecurity benefits of genetic engineering attribution. Nat Commun. 2020;11(1):9–12. doi:10.1038/s41467-020-19149-2

Li B, Si H-R, Zhu Y, et al. Discovery of Bat Coronaviruses through Surveillance and Probe Capture-Based Next-Generation Sequencing. mSphere. 2020;5(1):1–10. doi:10.1128/msphere.00807-19

Li D, Liu CM, Luo R, Sadakane K, Lam TW. MEGAHIT: An ultra-fast single-node solution for large and complex metagenomics assembly via succinct de Bruijn graph. Bioinformatics. 2015;31(10):1674–1676. doi:10.1093/bioinformatics/btv033

Li H, Handsaker B, Wysoker A, et al. The Sequence Alignment/Map format and SAMtools. Bioinformatics. 2009;25(16):2078–2079. doi:10.1093/bioinformatics/btp352

Li H. Minimap2: pairwise alignment for nucleotide sequences. Birol I, ed. Bioinformatics. 2018;34(18):3094–3100. doi:10.1093/bioinformatics/bty191

Liu S-L, Saif LJ, Weiss SR, Su L. No credible evidence supporting claims of the laboratory engineering of SARS-CoV-2. Emerg Microbes Infect. 2020;9(1):505–507. doi:10.1080/22221751.2020.1733440

Lu G, Wang Q, Gao GF. Bat-to-human: spike features determining ‘host jump’ of coronaviruses SARS-CoV, MERS-CoV, and beyond. Trends Microbiol. 2015;23(8):468–478. doi:10.1016/j.tim.2015.06.003

Luo C-M, Wang N, Yang X-L, et al. Discovery of Novel Bat Coronaviruses in South China That Use the Same Receptor as Middle East Respiratory Syndrome Coronavirus. J Virol. 2018;92(13):1–15. doi:10.1128/jvi.00116-18

Lusk RW. Diverse and Widespread Contamination Evident in the Unmapped Depths of High Throughput Sequencing Data. Gilbert T, ed. PLoS One. 2014;9(10):e110808. doi:10.1371/journal.pone.0110808

Martin D, Rybicki E. RDP: detection of recombination amongst aligned sequences. Bioinformatics. 2000;16(6):562–563. doi:10.1093/bioinformatics/16.6.562

Martin DP, Murrell B, Golden M, Khoosal A, Muhire B. RDP4: Detection and analysis of recombination patterns in virus genomes. Virus Evol. 2015;1(1):1–5. doi:10.1093/ve/vev003

Maynard Smith J. Analyzing the mosaic structure of genes. J Mol Evol. 1992;34(2):126–129. doi:10.1007/BF00182389

Meleshko D, Hajirasouliha I, Korobeynikov A. coronaSPAdes: from biosynthetic gene clusters to RNA viral assemblies. Robinson P, ed. Bioinformatics. 2021;38(1):1–8. doi:10.1093/bioinformatics/btab597

Menachery VD, Yount BL, Debbink K, et al. A SARS-like cluster of circulating bat coronaviruses shows potential for human emergence. Nat Med. 2015;21(12):1508–1513. doi:10.1038/nm.3985

Millet JK, Whittaker GR. Host cell entry of Middle East respiratory syndrome coronavirus after two-step, furin-mediated activation of the spike protein. Proc Natl Acad Sci. 2014;111(42):15214–15219. doi:10.1073/pnas.1407087111

Padidam M, Sawyer S, Fauquet CM. Possible Emergence of New Geminiviruses by Frequent Recombination. Virology. 1999;265(2):218–225. doi:10.1006/viro.1999.0056

Posada D, Crandall KA. Evaluation of methods for detecting recombination from DNA sequences: Computer simulations. Proc Natl Acad Sci. 2001;98(24):13757–13762. doi:10.1073/pnas.241370698

Prjibelski A, Antipov D, Meleshko D, Lapidus A, Korobeynikov A. Using SPAdes De Novo Assembler. Curr Protoc Bioinforma. 2020;70(1):1–29. doi:10.1002/cpbi.102

PyMOL. The PyMOL Molecular Graphics System, Version 2.0 Schrödinger, LLC.

Quay SC, Zhang D, Jones A, Deigin Y. Nipah virus vector sequences in COVID-19 patient samples sequenced by the Wuhan Institute of Virology. Published online September 19, 2021. http://arxiv.org/abs/2109.09112

Raj VS, Mou H, Smits SL, et al. Dipeptidyl peptidase 4 is a functional receptor for the emerging human coronavirus-EMC. Nature. 2013;495(7440):251–254. doi:10.1038/nature12005

Salminen MO, Carr JK, Burke DS, McCutchan FE. Identification of Breakpoints in Intergenotypic Recombinants of HIV Type 1 by Bootscanning. AIDS Res Hum Retroviruses. 1995;11(11):1423–1425. doi:10.1089/aid.1995.11.1423

Selitsky SR, Marron D, Hollern D, et al. Virus expression detection reveals RNA-sequencing contamination in TCGA. BMC Genomics. 2020;21(1):1–11. doi:10.1186/s12864-020-6483-6

Shinomiya N, Minari J, Yoshizawa G, Dando M, Shang L. Reconsidering the need for gain-of-function research on enhanced potential pandemic pathogens in the post-COVID-19 era. Front Bioeng Biotechnol. 2022;10(August):1–14. doi:10.3389/fbioe.2022.966586 SnapGene® software (from Dotmatics; available at snapgene.com).

Sun Y, Zhang H, Shi J, Zhang Z, Gong R. Identification of a Novel Inhibitor against Middle East Respiratory Syndrome Coronavirus. Viruses. 2017;9(9):255. doi:10.3390/v9090255

Tamura K, Stecher G, Kumar S. MEGA11: Molecular Evolutionary Genetics Analysis Version 11. Battistuzzi FU, ed. Mol Biol Evol. 2021;38(7):3022–3027. doi:10.1093/molbev/msab120

Tang XC, Zhang JX, Zhang SY, et al. Prevalence and Genetic Diversity of Coronaviruses in Bats from China. J Virol. 2006;80(15):7481–7490. doi:10.1128/JVI.00697-06

Teramichi T, Fukushi S, Hachiya Y, Melaku SK, Oguma K, Sentsui H. Evaluation of serological assays available in a biosafety level 2 laboratory and their application for survey of Middle East respiratory syndrome coronavirus among livestock in Ethiopia. J Vet Med Sci. 2019;81(12):1887–1891. doi:10.1292/jvms.19-0436

Thorvaldsdóttir H, Robinson JT, Mesirov JP. Integrative Genomics Viewer (IGV): High-performance genomics data visualization and exploration. Brief Bioinform. 2013;14(2):178–192. doi:10.1093/bib/bbs017

Vasimuddin M, Misra S, Li H, Aluru S. Efficient Architecture-Aware Acceleration of BWA-MEM for Multicore Systems. In: 2019 IEEE International Parallel and Distributed Processing Symposium (IPDPS). IEEE; 2019:314–324. doi:10.1109/IPDPS.2019.00041

Wang J-M, Wang L-F, Shi Z-L. Construction of a non-infectious SARS coronavirus replicon for application in drug screening and analysis of viral protein function. Biochem Biophys Res Commun. 2008;374(1):138–142. doi:10.1016/j.bbrc.2008.06.129

Wang Q, Qi J, Yuan Y, et al. Bat origins of MERS-CoV supported by bat Coronavirus HKU4 usage of human receptor CD26. Cell Host Microbe. 2014;16(3):328–337. doi:10.1016/j.chom.2014.08.009

Wang W, Peng X, Jin Y, Pan JA, Guo D. Reverse genetics systems for SARS-CoV-2. J Med Virol. 2022;94(7):3017–3031. doi:10.1002/jmv.27738

Waterhouse A, Bertoni M, Bienert S, et al. SWISS-MODEL: homology modelling of protein structures and complexes. Nucleic Acids Res. 2018;46(W1):W296–W303. doi:10.1093/nar/gky427

World Health Organization. Middle East respiratory syndrome coronavirus (MERS-CoV) – Saudi Arabia. 16 November 2022. https://www.who.int/emergencies/disease-outbreak-news/item/2022-DON422. Accessed 02/03/2023.

Woo PCY, Lau SKP, Li KSM, et al. Molecular diversity of coronaviruses in bats. Virology. 2006;351(1):180–187. doi:10.1016/j.virol.2006.02.041

Wu A, Wang Y, Zeng C, et al. Prediction and biochemical analysis of putative cleavage sites of the 3C-like protease of Middle East respiratory syndrome coronavirus. Virus Res. 2015;208:56–65. doi:10.1016/j.virusres.2015.05.018

Wu Z, Yang L, Ren X, et al. Deciphering the bat virome catalog to better understand the ecological diversity of bat viruses and the bat origin of emerging infectious diseases. ISME J. 2016;10(3):609–620. doi:10.1038/ismej.2015.138

Xia S, Lan Q, Pu J, et al. Potent MERS-CoV Fusion Inhibitory Peptides Identified from HR2 Domain in Spike Protein of Bat Coronavirus HKU4. Viruses. 2019;11(1):56. doi:10.3390/v11010056

Xie X, Muruato A, Lokugamage KG, et al. An Infectious cDNA Clone of SARS-CoV-2. Cell Host Microbe. 2020;27(5):841–848.e3. doi:10.1016/j.chom.2020.04.004

Xue LC, Rodrigues JP, Kastritis PL, Bonvin AM, Vangone A. PRODIGY: a web server for predicting the binding affinity of protein–protein complexes. Bioinformatics. 2016;32(23):3676–3678. doi:10.1093/bioinformatics/btw514

Yamada Y, Liu DX. Proteolytic Activation of the Spike Protein at a Novel RRRR/S Motif Is Implicated in Furin-Dependent Entry, Syncytium Formation, and Infectivity of Coronavirus Infectious Bronchitis Virus in Cultured Cells. J Virol. 2009;83(17):8744–8758. doi:10.1128/jvi.00613-09

Yang Y, Du L, Liu C, et al. Receptor usage and cell entry of bat coronavirus HKU4 provide insight into bat-to-human transmission of MERS coronavirus. Proc Natl Acad Sci. 2014;111(34):12516–12521. doi:10.1073/pnas.1405889111

Yang Y, Liu C, Du L, et al. (2015) Two Mutations Were Critical for Bat-to-Human Transmission of Middle East Respiratory Syndrome Coronavirus. J Virol. 2015;89(17):9119–9123. doi:10.1128/jvi.01279-15

Yount B, Denison MR, Weiss SR, Baric RS. Systematic Assembly of a Full-Length Infectious cDNA of Mouse Hepatitis Virus Strain A59. J Virol. 2002;76(21):11065–11078. doi:10.1128/JVI.76.21.11065-11078.2002

Yu P, Hu B, Shi Z-L, Cui J. Geographical structure of bat SARS-related coronaviruses. Infect Genet Evol. 2019;69(February):224–229. doi:10.1016/j.meegid.2019.02.001

Yuan Y, Cao D, Zhang Y, et al. Cryo-EM structures of MERS-CoV and SARS-CoV spike glycoproteins reveal the dynamic receptor binding domains. Nat Commun. 2017;8(1):15092. doi:10.1038/ncomms15092

Zhang W, Zheng X-S, Agwanda B, et al. Serological evidence of MERS-CoV and HKU8-related CoV co-infection in Kenyan camels. Emerg Microbes Infect. 2019;8(1):1528–1534. doi:10.1080/22221751.2019.1679610

